# Thiophenesulfonamides are specific inhibitors of quorum sensing in pathogenic Vibrios

**DOI:** 10.1101/2021.04.14.439868

**Authors:** Jane D. Newman, Priyanka Shah, Jay Chopra, Eda Shi, Molly E. McFadden, Rachel E. Horness, Laura C. Brown, Julia C. van Kessel

**Affiliations:** Department of Biology, Indiana University, Bloomington, IN 47405, USA; Indiana University School of Medicine, Indianapolis, IN 46202, USA; California Institute of Technology, Pasadena, CA 91125, USA; Department of Chemistry, Indiana University, Bloomington, IN 47405, USA

**Author notes:** <u>Corresponding authors:</u> Julia van Kessel, Laura Brown.

## Abstract

*Vibrio* bacteria are pathogens of fish, shellfish, coral, and humans due to contaminated seafood consumption. *Vibrio* virulence factors are controlled by the cell-to-cell communication called quorum sensing, thus this signaling system is a promising target for therapeutic design. We screened a compound library and identified nine compounds, including several 2-thiophenesulfonamides, that inhibit the master quorum sensing transcription factor LuxR in *Vibrio campbellii* but do not affect cell growth. We synthesized a panel of 50 thiophenesulfonamide compounds to examine the structure-activity relationship effects on quorum sensing *in vivo.* The most potent molecule identified, PTSP (3-phenyl-1-(thiophen-2-ylsulfonyl)-1*H*-pyrazole), specifically inhibits LuxR homologs in multiple strains of *Vibrio vulnificus*, *Vibrio parahaemolyticus*, and *V. campbellii* with sub-micromolar concentrations. PTSP efficacy is driven by amino acid conservation in the binding pocket, which is accurately predicted using *in silico* modeling of inhibitors. Our results underscore the potential for developing thiophenesulfonamides as specific quorum sensing-directed treatments for *Vibrio* infections.

## Introduction

*Vibrio* species are principal pathogens of marine animals, including fish, shellfish, and coral. Global warming and the concomitant rise of ocean temperatures correlates with increases in *Vibrio* prevalence and spread to regions beyond their typical equatorial habitats^1–3^, consequently harming fish and shellfish aquaculture industries and natural marine ecosystems worldwide. Thus, there is a global need for new treatments for vibriosis in coral reef ecosystems, aquaculture, and in human health, due to consumption of contaminated fish and shellfish. In pathogenic marine *Vibrio* species studied to date, the bacterial cell-cell signaling system called quorum sensing controls biofilm formation, as well as expression and secretion of virulence factors^4, 5^. Quorum sensing involves the production and detection of signaling molecules called autoinducers that provide information about the number and type of bacterial cells in the near vicinity. As populations of cells grow more dense, autoinducer concentrations increase, and detection of these molecules drives changes in gene expression to alter population-wide behaviors, including those required for pathogenesis.

In vibrios, autoinducers are sensed by membrane-bound histidine kinase receptors that participate in a phosphorylation cascade, ultimately controlling production of the master regulator LuxR (Fig. 1)^5–7^. At low cell densities (LCD), low levels of LuxR are produced. At high cell densities (HCD), maximal LuxR protein is produced, and this transcription factor activates and represses hundreds of genes^8^. Although the number and type of autoinducers and receptors vary among vibrios, the LuxR protein is highly conserved in all pathogenic vibrios studied to date^9–14^. Although the naming of the *V. campbellii* protein LuxR causes confusion, this protein does not resemble or function like the LuxR protein that is part of the *Vibrio fischeri* LuxI/LuxR quorum sensing system, which requires binding to the autoinducer molecule made by LuxI for activity^15^. Conversely, the LuxR from *V. campbellii* belongs to the TetR superfamily^16^, and these proteins are structurally, genetically, biochemically, and functionally distinct from the *V. fischeri* LuxR. LuxR/TetR homologs in vibrios include SmcR in *V. vulnificus*, HapR in *V. cholerae*, OpaR in *V. parahaemolyticus*, and VcpR in *Vibrio coralliilyticus*, which share 76-96% amino acid identity^14^. The LuxR/TetR-type proteins do not have a known ligand, although a putative ligand binding pocket has been defined in structures and shown to bind inhibitors^17–19^. LuxR/TetR proteins in vibrios directly bind to multiple sites in promoter regions and interact with other proteins *(e.g.*, RNA polymerase, IHF) or compete with other proteins *(e.g.*, H-NS) to activate or repress transcription of hundreds of quorum sensing genes^6, 8, 20–22^.

**Figure 1.**
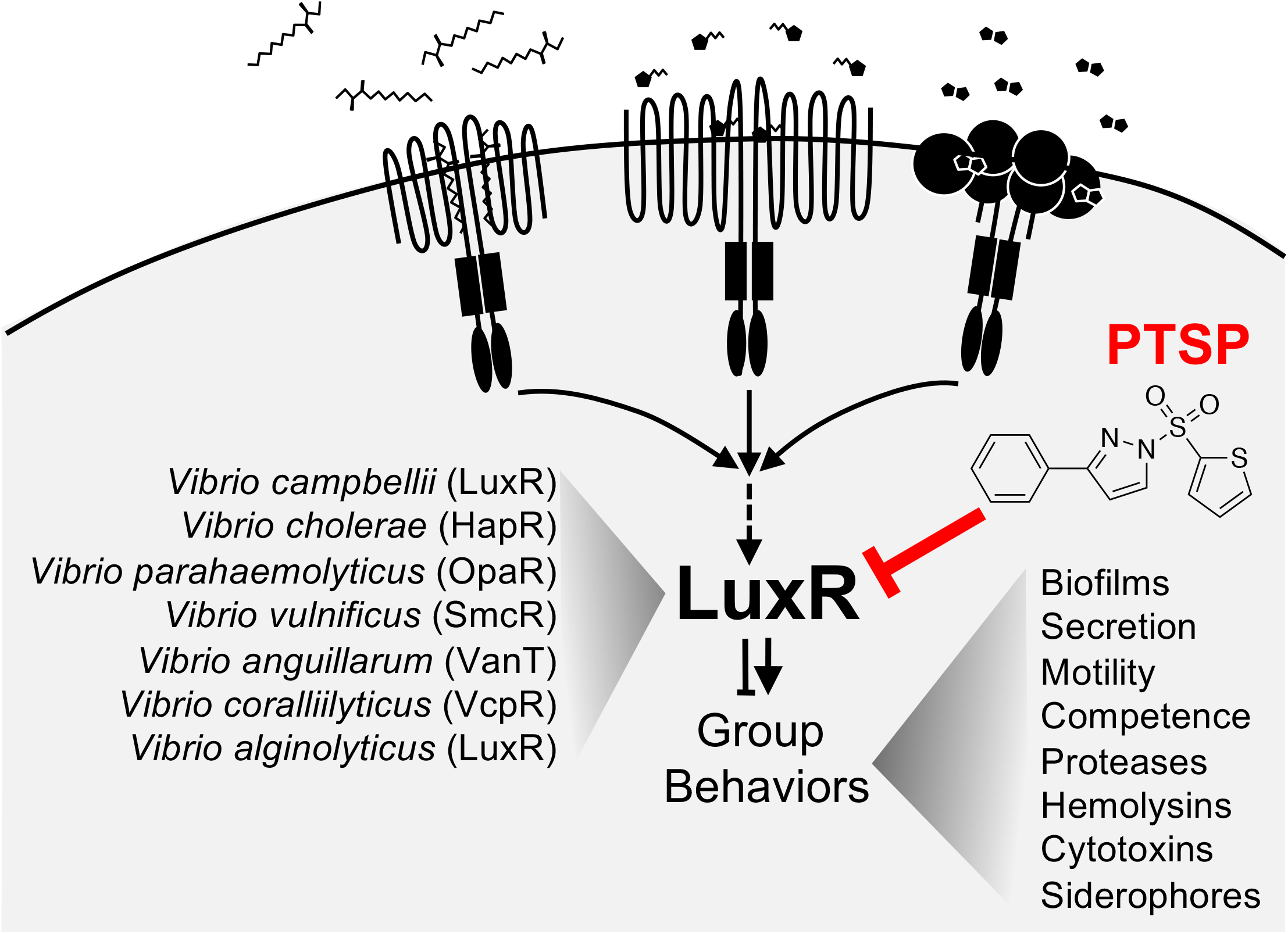
The quorum sensing pathway in *Vibrio* species. Autoinducer molecules are bound by membrane-bound histidine kinase receptors, which alters the phosphorylation cascade downstream. At high cell densities, the production of the transcription factor LuxR is maximal, and LuxR activates and represses genes encoding proteins with various functions, some of which are listed in the diagram. LuxR protein homologs in various *Vibrio* species are listed.

*Vibrio* LuxR/TetR-type proteins play crucial roles in colonization and infection of hosts through quorum-directed regulation of biofilm formation, type III and type VI secretion systems, motility, and production of proteases, hemolysins, siderophores, and cytotoxins^4^. Deletion or inhibition of LuxR proteins in several vibrios reduces or eliminates colonization and toxicity, effectively increasing host survival^11,17,23^. Thus, LuxR represents a key target for designing therapeutics to block quorum sensing in vibrios. LuxR inhibitors would presumably render *Vibrio* cells unresponsive to quorum sensing signals even at HCD, thus restricting cells to their LCD gene expression program. Indeed, recent studies have shown that quorum sensing inhibitors are viable alternative approaches to traditional antibiotics in disease treatment with demonstrated efficacy in animal models and active clinical trials^24,25^. Because quorum sensing inhibitors typically do not affect growth of the bacteria but rather inhibit specific pathways^26,27^, these molecules are hypothesized to generate less selective pressure for evolving resistance.

*V. campbellii* LuxR-specific inhibitors have been identified via a variety of methods, including both *in vitro* and *in silico* screening strategies. These include compounds with varying functional groups such as aromatic enones, sulfonamides, sulfones, cinnamaldehydes, furanones, and brominated thiophenones^28–33^, and each was shown to specifically inhibit bioluminescence, biofilm formation, and/or protease activity in *V. campbellii* and in some cases other *Vibrio* species. However, the range of inhibition for some of these molecules is either low (~3-fold) or the inhibitory concentrations required to observe phenotypic effects *in vitro* or *in vivo* are high (>20 μM). Several of these molecules show favorable therapeutic potential because addition of these molecules to brine shrimp larvae infected with *Vibrio* cells increases survival. In particular, the molecule called Qstatin (1-((5-bromothiophen-2-yl)sulfonyl)-1*H*-pyrazole) is a highly promising molecule that was shown to be a specific inhibitor of the LuxR homolog SmcR in *Vibrio vulnificus* both *in vitro* and *in vivo*^17^.

Here, we used an *E. coli* bioassay to screen chemical libraries to identify specific inhibitors of LuxR that do not affect bacterial cell growth. Our structure-activity relationship data shows that multiple thoiphenesulfonamide-containing molecules with heterocycle variations are strong inhibitors of LuxR-type proteins in a wide-range of pathogenic *Vibrio* species.

## Results

### A bioassay screen identifies LuxR chemical inhibitors

LuxR proteins activate and repress gene expression in vibrios through direct binding to specific DNA sequences in promoters^6, 14, 22^. Previously, we developed a bioassay that consists of a dual-color fluorescent reporter plasmid that reports both LuxR activities: the *luxCDABE* promoter is activated by LuxR and drives expression of *gfp*, and the *VIBHAR_05222* promoter is repressed by LuxR and drives expression of *mCherry*^22^ (Fig. 2A). We use this reporter plasmid in an *E. coli* strain that also contains a plasmid expressing LuxR from its native promoter. Thus, in the presence of LuxR, GFP levels increase and mCherry levels decrease compared to the control strain (Fig. 2B). As a positive control, we showed that Qstatin inhibits LuxR activity in this assay (Fig. S1). We screened ~60,000 molecules in the ChemBridge (Fig. S1A, S1C) and Chemdiv (Fig. S1B) libraries and identified nine compounds that inhibit LuxR activation and/or repression but do not affect the final growth yield more than 10%. Four of these compounds contain a sulfonamide or sulfamide core with variable groups on each side similar to Qstatin, whereas the other compounds are structurally dissimilar (Fig. 2C). We synthesized or purchased these molecules and determined the IC_50_ for each in the *E. coli* bioassay strain using titration curves (Table S1, Fig. 2D, 2E). P0053 I18 has the best inhibitory effect on LuxR in the *E. coli* bioassay with an IC_50_ similar to Qstatin (Table S1, Fig. 2D, 2E). P2065 E16 has only a minor inhibition of LuxR, and a variation of this molecule lacking the CF3 group on the heterocycle has no activity (Fig. 2D, 2E). We also observed consistent though low inhibition by P0074 H04 and P0053 O05, which also contain sulfamide/sulfonamide cores (Fig. 2D, 2E).

**Figure 2.**
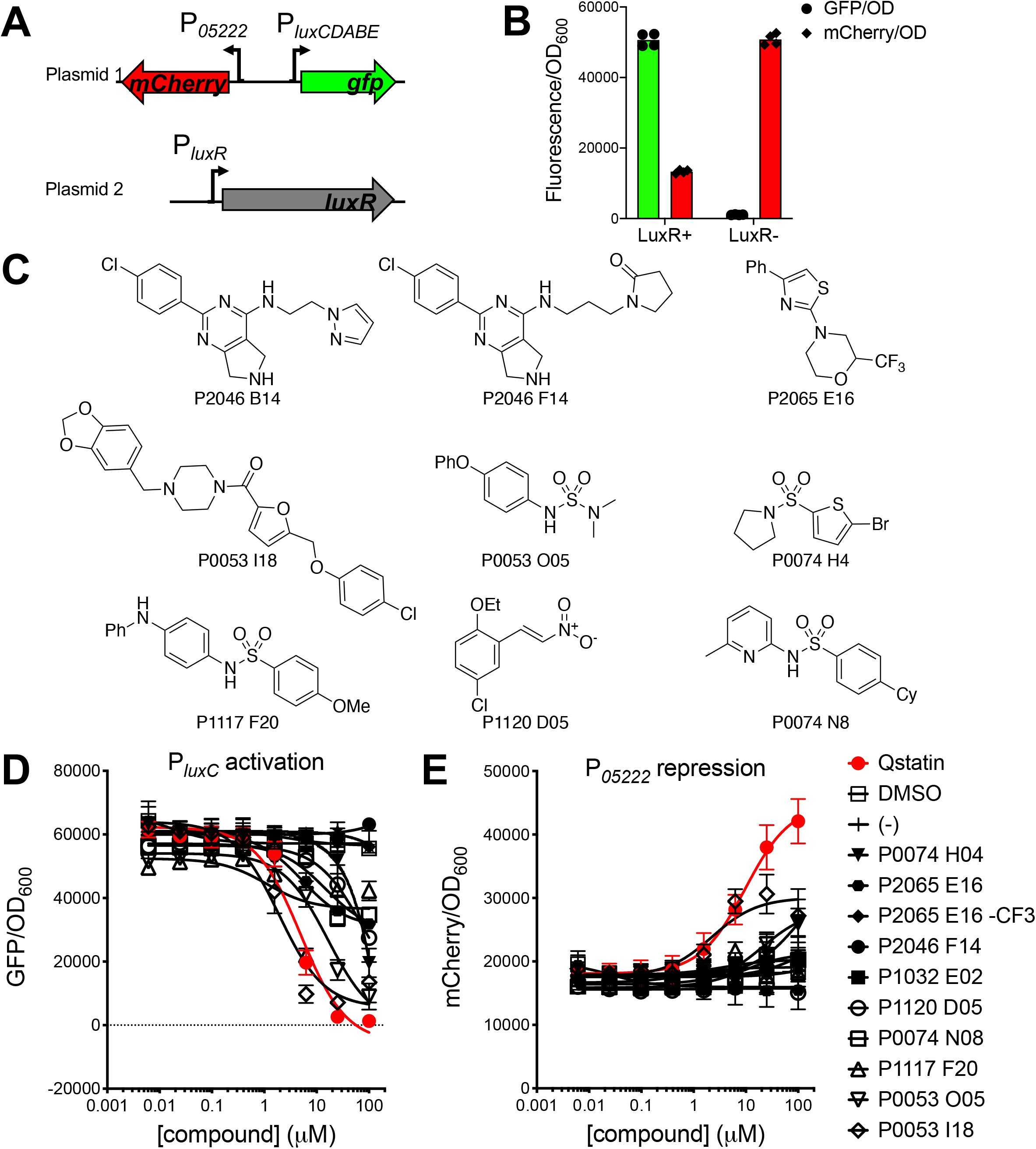
Screen for LuxR inhibitors in *E. coli* bioassay. (A) Diagram of the two plasmids in the *E. coli* bioassay used to screen for LuxR inhibitors. Plasmid 1 (pJV064) contains divergent promoters for *V. campbellii luxCDABE* and *05222* driving expression of *gfp* and *mCherry*, respectively. Plasmid 2 (pKM699) contains the *V. campbellii luxR* gene under control of its native promoter. (B) Fluorescence expression (GFP/OD_600_ or mCherry/OD_600_) in *E. coli* bioassay cells (with pJV064) expressing LuxR (pKM699) or the empty vector control (pLAFR2). (C) Structures of LuxR inhibitors identified and verified from the compound library screen. (D, E) Production of GFP (panel D; GFP/OD_600_) or mCherry (panel E; mCherry/OD_600_) in the presence of LuxR inhibitors titrated into the *E. coli* bioassay strain (pKM699, pJV064). DMSO was titrated as a solvent control with an equal volume to the 100 μM concentration of compound and compared to cells in which nothing was added (-, plotted at 100 μM point on x-axis). Data shown represent the mean and standard deviation of at least three biological replicates.

### 2-thiophenesulfonamide compounds specifically inhibit LuxR

To further explore the thiophenesulfonamide class of compounds that includes Qstatin and P0074 H4, we synthesized a panel of Qstatin derivatives with steric and electronic structural variations in both heterocycles (Fig. 3A). We refer to modifications to the heteroaromatic amine ring (pyrazole in the case of Qstatin) with number designations for each class and modifications to the thiophene ring with letter designations for each class (Fig. 3A). Assays with these molecules in the *E. coli* bioassay showed that the most active compounds contain a 3-methyl- (class 8) or 3-phenyl-substituted pyrazole (class 10), an unsubstituted pyrazole (class 1), or a pyrrole in place of the pyrazole (class 3) (Fig. 3B, 3C, S2). Compounds containing other heterocycles do not have activity against LuxR (Fig. S2). It is particularly noteworthy that the compounds containing an imidazole ring (class 2) are not active given that the structure is highly similar to pyrroles and pyrazoles (classes 1 and 3). Methyl substitution at the 3 position of the pyrazole does not alter activity compared to Qstatin (class 8), however methyl substitution at both the 3- and 5-positions of the pyrazole eliminates activity (class 9). In most cases, the presence/absence of Br or Cl atoms on the thiophene ring does not alter activity (Fig. 3). For example, Qstatin, 1B, and 1C have similar activities, and 10A, 10B, and 10C have similar activities. The conformation of the sulfonamide core appears to be critical because substitution with a carbonyl eliminated activity of that class of compounds (classes F and G). The most potent molecules are compounds 10A, 10B, and 10C, all of which contain a phenyl group on the 3-position of the pyrazole and vary in the presence or absence of bromine or chlorine on the thiophene ring (Fig. 3, S2).

**Figure 3.**
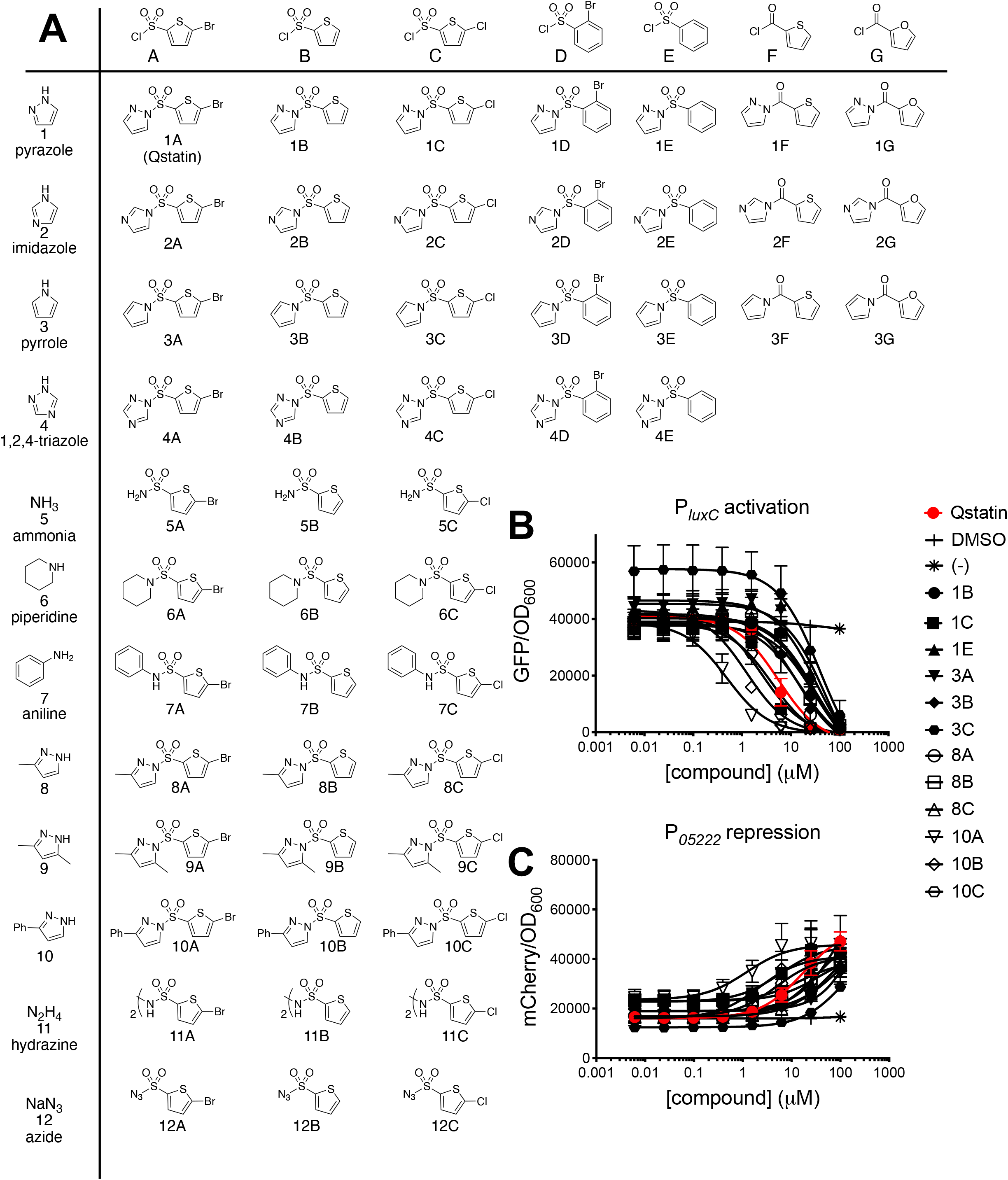
Panel of thiophenesulfonamide molecules that inhibit LuxR. Substrates with modifications to the heteroaromatic ring (pyrazole in the case of Qstatin) have number designations, and substrates with modifications to the thiophene ring have letter designations. (B, C) Production of GFP (panel B; GFP/OD_600_) or mCherry (panel C; mCherry/OD_600_) in the presence of thiophenesulfonamide compounds titrated into the *E. coli* bioassay strain (pKM699, pJV064). DMSO was titrated as a solvent control with an equal volume to the 100 μM concentration of compound and compared to cells in which nothing was added (-, plotted at 100 μM point on x-axis). Data shown represent the mean and standard deviation of at least three biological replicates.

Many sulfonamide-containing compounds are known to target the folate synthesis pathway in bacteria and inhibit growth (bacteriostatic), and thus these “sulfa drugs” have been used as broad-spectrum antibiotics for decades^34,35^. However, the structurally distinct 2-thiophenesulfonamides generally do not have any bacteriostatic activity; only two compounds (4A and 4C) in this entire panel limit *E. coli* growth yield more than 10% compared to negative controls, and this effect was only observed at concentrations of 100 μM and higher (Fig. S3A). Addition of high concentrations of 10B or 10C to *V. campbellii* or *V. vulnificus* does not alter growth rate or growth yield (Fig. S3B, 3C), suggesting that the inhibitory activity is specific to LuxR, and these are not general antibacterial compounds. Indeed, RNA-seq performed with Qstatin showed a specific effect on the SmcR regulon^17^. Further, we note that molecule 10B inhibits LuxR activity rapidly; GFP production is blocked 120 minutes after 10B addition to the culture, and this is maintained until 16 hours (Fig. S3D). Collectively, these data show that 2-thiophenesulfonamide molecules are stable in bacterial culture, specifically inhibit LuxR, and do not significantly affect cell growth.

### 2-thiophenesulfonamides inhibit quorum sensing in Vibrios

Qstatin has been shown to be an effective inhibitor of SmcR *in vitro* and *in vivo*^17^. It also inhibits pathogenesis in *V. campbellii, V. parahaemolyticus*, and *V. vulnificus* in a shrimp infection assay, presumably through inhibition of the LuxR-type protein in these strains^17^. To assess the activity of our panel of thiophenesulfonamide inhibitors against other vibrios, we assayed quorum-sensing controlled phenotypes in several strains from five *Vibrio* species: *V. campbellii, V. coralliilyticus, V. cholerae, V. parahaemolyticus*, and *V. vulnificus.* Although most of these vibrios are not bioluminescent, the *luxCDABE* operon from *V. campbellii* BB120 is routinely used to measure quorum sensing and LuxR regulation in other vibrios because LuxR directly binds this promoter and is required for gene expression^17,36–39^. We introduced a plasmid containing the *luxCDABE* promoter driving expression of *gfp* (pCS19 (kanamycin-resistance cassette) or pCS42 (gentamicin-resistance cassette)) into each of the *Vibrio* strains. We focused our assays on the molecules with the most activity determined in Fig. 3, which we refer to as the “top panel”. We observed that 10A, 10B, and 10C molecules are the most inhibitory in *V. campbellii, V. vulnificus*, and *V. parahaemolyticus*, and in each, 8A has a similar IC_50_ to Qstatin (Fig. 4A-E, Table S2). Molecules 10A, 10B, and 10C are so potent in *V. vulnificus* that we performed extended serial dilutions to obtain accurate IC_50_ data because initial titrations did not yield enough points for a complete curve (Fig. 4E, S4A-C, Table S2). We also observed that few molecules are active in *V. coralliilyticus* OCN008, but 3B has the most noticeable effect (Fig. 4C). Importantly, there is a distinct difference in the half-maximal inhibitory concentration (IC_50_) for each molecule when compared across the five species. *V. vulnificus* exhibits very low IC_50_ values for all the top panel molecules (Fig. 4E), whereas *V. cholerae* is completely resistant to these molecules even at high concentrations (Fig. 4B). Using 10B as an example, the IC_50_ values are orders of magnitude different comparing *V. vulnificus* (0.002 μM) to *V. campbellii* (0.35 μM) (Fig. 4A, 4E, Table S2). To further examine this observation, we assayed the top panel of molecules against additional isolates for each species (Table S2). We observed that the inhibitory effect of the molecules is similar for each isolate within a species (Table S2). For example, the IC_50_ values for 10B for all three *V. vulnificus* isolates range from 2-30 nM. From these data, we conclude that 10B has the highest inhibitory activity in all *Vibrio* species, with variation in the IC_50_ that is specific to the *Vibrio* species tested.

**Figure 4.**
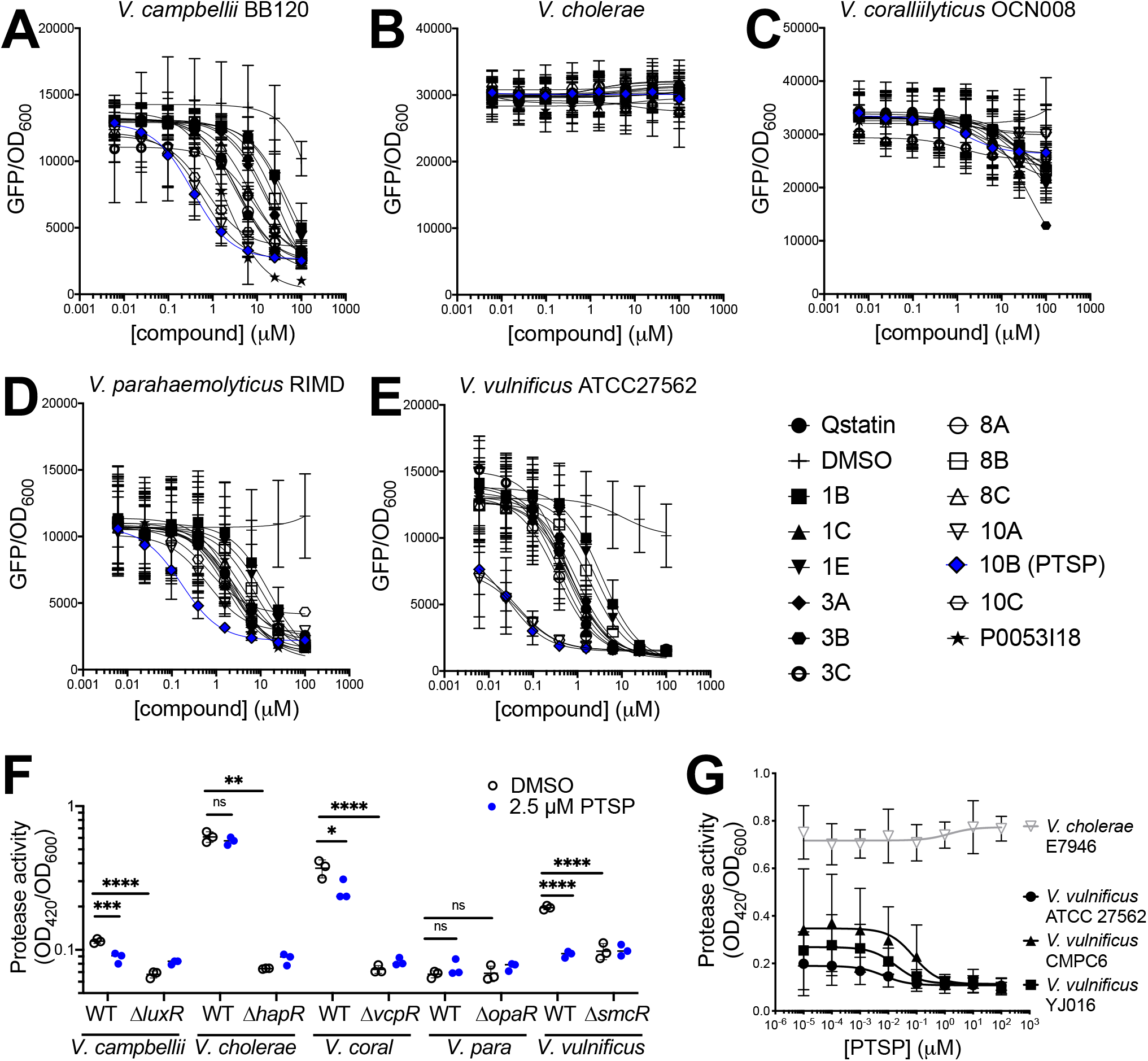
Thiophenesulfonamides have a range of inhibition against *Vibrio* species. (A-E) Titration of molecules from the top panel of thiophenesulfonamides in *Vibrio* strains compared to DMSO solvent control (DMSO was titrated with an equal volume to the 100 μM concentration of compound). Data shown represent the mean and standard deviation of three biological replicates. (F) Protease activity (final assay OD_420_/initial culture OD_600_) for wild-type and mutant *Vibrio* strains in the presence of 2.5 μM PTSP or an equal volume of the DMSO solvent. Asterisks indicate significant differences (two-way analysis of variance (ANOVA) followed by Sidak’s multiple comparisons test, *n* = 4; *, *p* = 0.05; **, *p* = 0.01; ***, *p* = 0.001; ****, *p* = 0.0001; ns, not significant). (G) Protease activity (final assay OD_420_/initial culture OD_600_) for *Vibrio* strains in the presence of PTSP titrated into the cultures. Data shown represent the mean and standard deviation of three biological replicates.

We next focused on the best inhibitor in the panel, 10B, which is called 3-<u>p</u>henyl-1- (<u>t</u>hiophen-2-yl<u>s</u>ulfonyl)-1*H*-<u>p</u>yrazole, and thus we will refer to it as PTSP from hereon. We assessed LuxR function in the presence of PTSP by assaying protease production, a key virulence activity in vibrios during pathogenesis and host cell lysis (Fig. 4F, 4G). LuxR proteins activate several genes encoding proteases in *Vibrio* species, such as HapA in *V. cholerae*, VvpE in *V. vulnificus*, PrtA in *V. parahaemolyticus*, and VcpA/VcpB in *V. coralliilyticus* (previously called VtpA and VtpB when the strain was misidentified as *Vibrio tubiashii*)^23,40–43^. We analyzed protease activity of bacterial supernatants using azocasein as a substrate and compared wild-type strains to their isogenic Δ*luxR* counterparts for one representative strain for each species. In each species except *V. parahaemolyticus*, the wild-type strain produces significantly more protease activity than the Δ*luxR* strain (Fig. 4F). Addition of PTSP to the wild-type strain significantly reduces protease activity in *V. campbellii, V. coralliilyticus*, and *V. vulnificus*, but not in *V. cholerae* (Fig. 4F). *V. parahaemolyticus* exhibited extremely low protease activity in this assay, thus the effect of PTSP on *V. parahaemolyticus* cannot be ascertained in this experiment. We observed that the effect of PTSP on protease activity mirrors that of bioluminescence: PTSP has minimal activity against *V. coralliilyticus* and highest activity against *V. vulnificus.* (Fig. 4F). Although we assessed the effect of PTSP on protease activity in numerous strains for *V. campbellii*, *V. coralliilyticus*, and *V. parahaemolyticus*, the protease activity is so low in many of these isolates that an IC_50_ could not be reliably calculated for these (Fig. S4). However, for *V. vulnificus* strains, we calculated the IC_50_ values for PTSP inhibition and observed a similar range of protease inhibition as we observed for bioluminescence: ATCC 27562 = 6.8 nM, CMPC6 = 78.1 nM, and YJ016 = 18.3 nM (Fig. 4G). From the bioluminescence and protease assay data, we conclude that PTSP is a potent inhibitor of *V. vulnificus, V. parahaemolyticus*, and *V. campbellii*, with moderate effects on *V. coralliilyticus.* Inhibitor PTSP and derivatives are not active against *V. cholerae* (Fig. 4B, 4F, 4G). Further, we conclude that the use of the *E. coli* bioassay reporter is a valid assay for monitoring endogenous LuxR activity in these *Vibrio* species.

### Mechanism of 2-thiophenesulfonamide efficacy

There are several possible reasons for the observed difference in PTSP inhibition in the various *Vibrio* species, including but not limited to: 1) differences in LuxR-inhibitor interaction(s), 2) differences in diffusion of the inhibitor across the membranes, or 3) stability of the inhibitor in the cell and/or cell culture. To examine PTSP activity in a common strain background, we cloned the *luxR* gene from each *Vibrio* species into a plasmid under control of an IPTG-inducible promoter and assayed the effect of PTSP against these proteins in the *E. coli* bioassay. We found that HapR is unresponsive to PTSP in *E. coli*, exhibiting a similar level of GFP expression to the negative control (Fig. 5B). Conversely, SmcR, LuxR, OpaR, and VcpR are each inhibited by PTSP with a trend similar to that observed in their native *Vibrio* cells, in which the order of sensitivity to PTSP inhibition is SmcR>LuxR>OpaR>VcpR (Fig. 5B). We conclude that the differences in PTSP activity in *Vibrio* species is due to differences in interaction between PTSP and the LuxR-type protein in each *Vibrio.*

**Figure 5.**
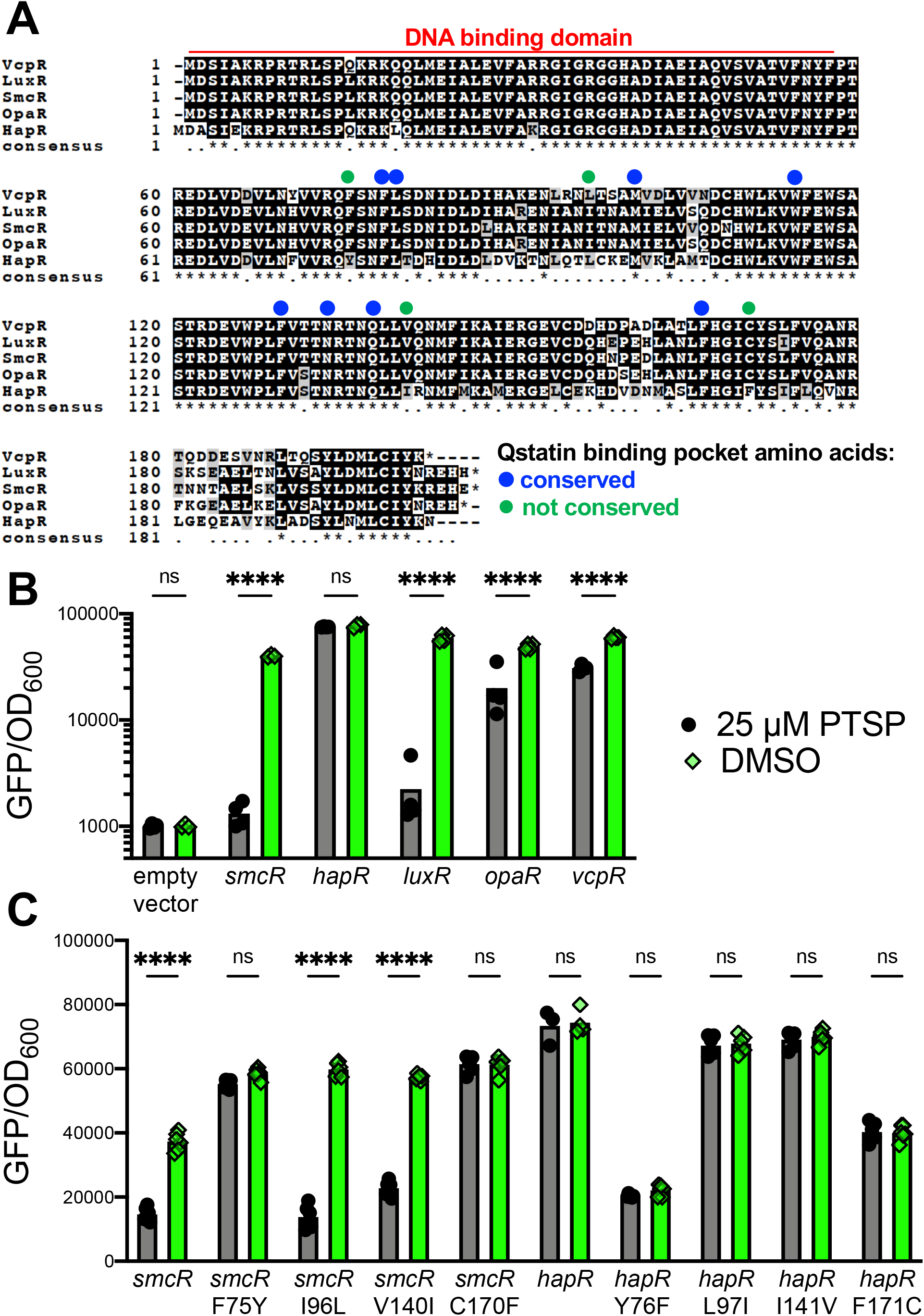
Sulfonamide 10B does not inhibit *V. cholerae* HapR. (A) LuxR sequences from *V. coralliilyticus* OCN008 (accession: ERB64458.1), *V. campbellii* BB120 (accession: ABU72404.1), *V. vulnificus* ATCC 27562 (accession: WP_011079558.1), *V. parahaemolyticus* RIMD 2210633 (accession: WP_005479697.1), and *V. cholerae* C6706 (accession: ACB30340.1) aligned using Clustal Omega and Boxshade. Blue dots indicate the amino acids in SmcR that contact Qstatin in the published SmcR-Qstatin X-ray crystal structure^17^. Green dots indicate amino acid differences between LuxR proteins in each species among the SmcR-Qstatin contacts. (B) Production of GFP (GFP/OD_600_) in the presence of 25 μM PTSP or DMSO solvent control in the *E. coli* bioassay strain (pJV064) containing plasmids expressing SmcR (pJN22), HapR (pJV387), LuxR (pJV388), OpaR (pJV389), VcpR (pJV390), or empty vector (pMMB67EH-kanR). SmcR and HapR expression in strains was induced with 50 μM IPTG; IPTG was not added to cultures of the remaining strains. (C) Production of GFP (GFP/OD_600_) in the presence of 25 μM PTSP or DMSO solvent control in the *E. coli* bioassay strain (pJV064) containing plasmids expressing SmcR (pJN22), HapR (pJV387), or the listed various amino acid substitution mutants. SmcR and HapR expression was induced with 50 μM IPTG. For panels B and C, asterisks indicate significant differences (two-way ANOVA followed by Sidak’s multiple comparisons test, *n* = 3 (panel B), *n* = 6 (panel C); ****, *p* = 0.0001; ns, not significant).

Using the SmcR-Qstatin X-ray crystal structure as a guide^17^, alignment of the LuxR-type proteins from each *Vibrio* species shows that there are four residues that interact with Qstatin in SmcR that are variable in HapR and/or VcpR (Fig. 5A). We therefore sought to determine if any of the four residues that differ between HapR and LuxR/SmcR in the putative ligand binding pocket are sufficient to render SmcR insensitive to PTSP. First, we introduced substitutions in SmcR to mimic the amino acid sequence of HapR: F75Y, I96L, V140I, and C170F. We observed that the substitutions of F75Y and C170F abolish PTSP inhibition, suggesting that F75 and C170 are both necessary for PTSP inhibition of SmcR transcription regulation *in vivo* (Fig. 5C). Next, we introduced single substitutions in HapR to mimic the amino acid present in SmcR: Y76F, L97I, I141V, and F171C. However, none of these substitutions alone are sufficient to make HapR sensitive to PTSP (Fig. 5C). From these results, we conclude that SmcR F75 and C170 are critical residues necessary for PTSP inhibition of SmcR activity.

### Modeling of 2-thiophenesulfonamide inhibitors

Previous studies have used molecular docking simulations to predict effective inhibitors of LuxR family proteins^30^. To examine the accuracy of using molecular docking to predict effective inhibitors of LuxR proteins, we used Autodock Vina software^44^ to simulate binding of molecules to the structure of SmcR and calculate the best binding mode and affinity (kcal/mol). First, to validate this modelling approach, we used Autodock Vina to predict the binding position of Qstatin into the SmcR (apo) X-ray crystal structure and compared it to the solved structure of SmcR-Qstatin (Fig. 6A). Although there is a slight shift (0.6 to 1.4 Å), the position and orientation of Qstatin within the putative ligand binding pocket of SmcR was accurately predicted by Autodock Vina (Fig. 6A). We next used Autodock Vina to predict the binding position of PTSP in both SmcR and HapR (Fig. 6B). We observed that PTSP is modeled in the opposite orientation in SmcR compared to HapR. This is likely driven at least partially by the rotational position of glutamine 137 in SmcR (Q138 in HapR) that clashes with the phenyl ring of PTSP. We also note that SmcR Q137 has multiple rotamers among the four SmcR chains within the asymmetric unit (Fig. 6C), and Autodock Vina modelling indicates that PTSP is oriented differently in SmcR chain A compared to chain B (Fig. 6D). Thus, the binding orientation and interactions of PTSP with SmcR are likely influenced, at least in part, by the orientation of Q137. For example, the predicted orientation of PTSP is different in chains A and B that have different Q137 rotamers and different predicted binding energies (Fig. S5; −7.5 kcal/mol for chain A compared to −0.2 kcal/mol for chain B). In addition, the predicted binding energies for PTSP in HapR are clearly worse at +1.6 kcal/mol for chain A and +10.4 kcal/mol for chain B (Fig. S5).

**Figure 6.**
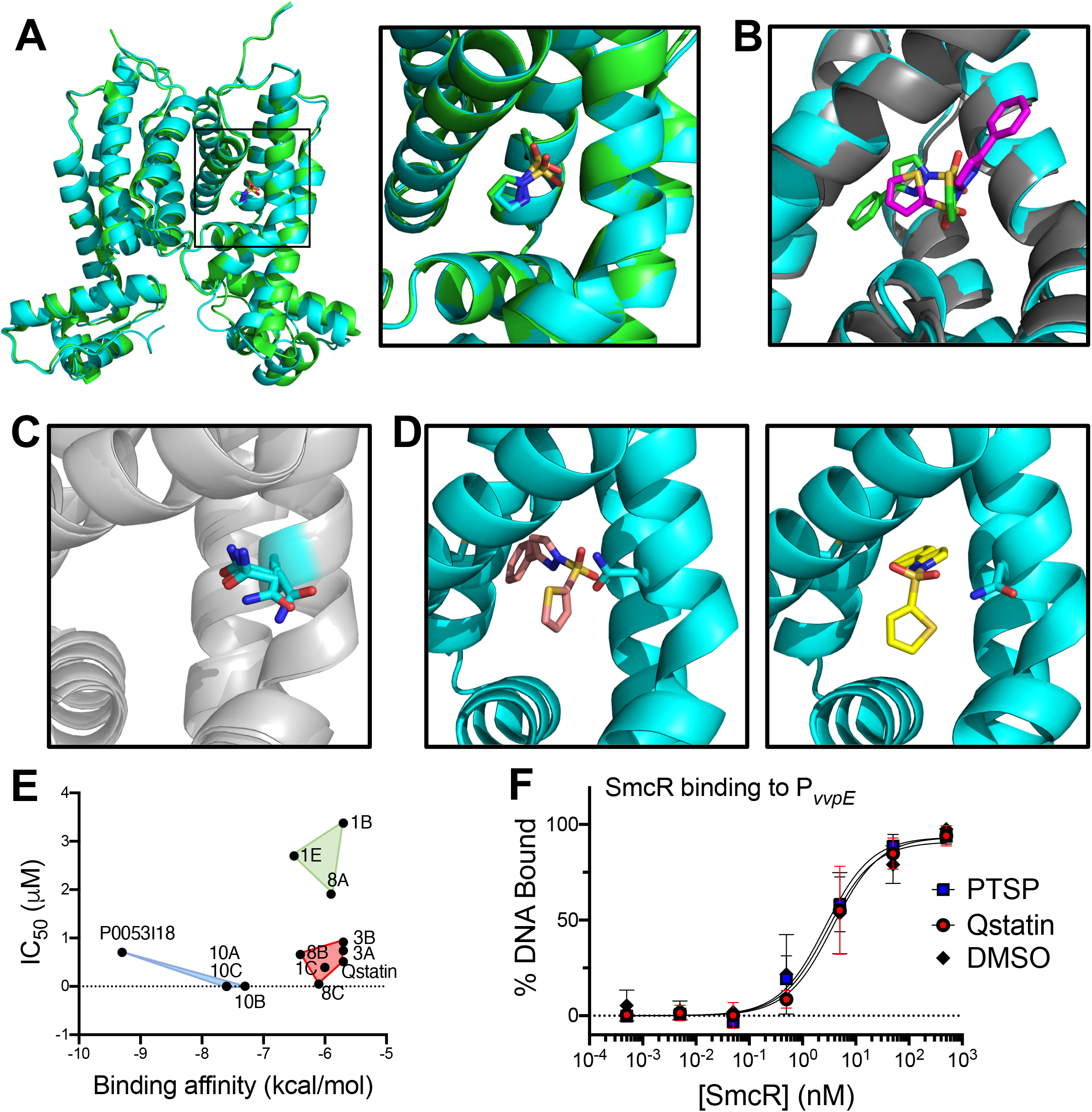
(A) Autodock Vina modelling of Qstatin into the SmcR apo X-ray crystal structure (3KZ9; cyan) overlaid with the X-ray structure of SmcR complexed with Qstatin (5X3R; green). (B) Autodock Vina modelling of PTSP (10B) in the X-ray crystal structures of SmcR (3KZ9; cyan) and HapR (2PBX; gray). (C) The rotational orientation of Q137 in the four chains of SmcR from the asymmetric unit (3KZ9). (D) Autodock Vina modelling of PTSP into chain A (left) and chain B (right) of SmcR (3KZ9). (E) Binding affinities predicted by Autodock Vina for each molecule modelled into the SmcR X-ray crystal structure (3KZ9) are graphed against the IC_50_ value for each molecule determined by assaying bioluminescence production in *V. vulnificus* ATCC27562 (Table S2). Clusters of the 4 (blue), 6 (red), and 3 (green) data points were generated using K-means clustering analysis in R. (F) Electrophoretic mobility shift assays (EMSAs) of SmcR purified protein in varying concentrations incubated with radiolabeled dsDNA corresponding to the *V. vulnificus vvpE* promoter. DNA shifts were quantified using ImageJ, and the graphs show the mean and standard deviation for three biological replicates.

Because Autodock Vina accurately predicts the binding position of Qstatin in SmcR, we modelled binding of the top panel of molecules into SmcR. We plotted the predicted binding affinities for each molecule against the calculated IC_50_ values from the *in vivo* assay in *V. vulnificus* (Fig. 6E). Molecules 10A, 10B (PTSP), and 10C cluster with P0053 I18 in a group with the lowest predicted binding energies and lowest observed IC_50_ values (Fig. 6E). Conversely, the molecules with the low-moderate inhibitory activity were predicted to have similar binding energies (Fig. 6E). However, these molecules group in two separate clusters, indicating that modelling could not distinguish efficacy among low-moderate inhibitors. From these data, we conclude that molecular docking simulations are an accurate method for predicting molecules with good binding affinity and inhibitory activity against SmcR.

We note that Qstatin and PTSP are both predicted to bind in the putative ligand binding pocket of SmcR. It is currently unknown how Qstatin allosterically affects DNA binding through its interactions in the ligand binding domain. Kim *et al.* showed that Qstatin does not appreciably alter the DNA binding constant for SmcR, but rather affects the entropy and enthalpy with which it interacts with DNA^17^. We therefore also assessed the effect of sulfonamide PTSP on the DNA binding activity of LuxR and SmcR *in vitro.* Addition of saturating concentrations of PTSP does not alter DNA binding to LuxR or SmcR (Fig. 6F, Fig. S6). This result is similar to the finding that Qstatin has very little effect on SmcR DNA binding affinity (Fig. 6F)^17^. Collectively, these data show that thiophenesulfonamides inhibit the function of LuxR proteins to different levels. In addition, these data suggest that the mechanism of inhibition by thiophenesulfonamides is likely similar in both SmcR and LuxR.

## Discussion

This study has aimed to test a panel of inhibitors against quorum sensing, a non-essential cell signaling pathway that controls pathogenesis in *Vibrio* species. Using a previously established dual-color bioassay, we screened thousands of compounds and synthesized a broad panel to find potent inhibitors of the quorum sensing master transcription factor LuxR^8,22^. We observed that some of our key candidates contain sulfamide/sulfonamide heterocycles, but there exists a vast range of inhibition between these various compounds. Using the best candidate inhibitor PTSP, we tested its efficacy against five vibrios *in vivo* and found that it is most effective towards *V. vulnificus*, followed by *V. campbellii, V. parahaemolyticus, V. coralliilyticus*, and not effective against *V. cholerae.* The efficacy observed in inhibition of bioluminescence reporter assays was mimicked in protease assays where measurable. These results show that PTSP blocks all measured activities of LuxR proteins. This result is comparable to what was observed for the thiophenesulfonamide Qstatin, which blocks SmcR regulation of genes across the *V. vulnificus* genome to similar levels as a Δ*smcR* strain^17^. Thus, our results show that thiophenesulfonamides are broadly inhibitory of LuxR activities in multiple *Vibrio* species.

We were intrigued by the finding that the compounds we tested have no effect on HapR from *V. cholerae.* Our data show that HapR resistance is due to amino acid residue differences in the putative ligand binding pocket. We focused on differences in the residues in the Qstatin binding pocket across five *Vibrio* species, though substitution of single amino acids in HapR are not sufficient to render the protein sensitive to PTSP. We note that substitutions Y76F and F171C did not alter sensitivity to the PTSP molecule, but rather affected overall activity of HapR. We presume that multiple, combined substitutions in the HapR ligand binding pocket to change the sequence to match SmcR would likely result in HapR sensitivity to PTSP, though this needs to be formally tested. Interestingly, *V. cholerae* also has a very different pathogenic life cycle compared to other vibrios. Pathogenesis in the human host caused by *V. cholerae* occurs at LCD, where the bacterial cells attach to intestinal epithelial cells through the toxin co-regulated pilus and grow as a biofilm, producing cholera toxin. Growth of the population and accumulation of autoinducers drives inhibition of biofilms through various regulatory mechanisms, cleavage from the host epithelium by the HapA protease, and the cells are shed back into the marine environment^45–49^. This poses some intriguing evolutionary questions about *V. cholerae* pathogenesis and growth in the environment and the selective pressures that may have driven differences in amino acid conservation between HapR and other LuxR-family proteins. This could underscore the stark contrast in efficacy of inhibitors against HapR and other *Vibrio* LuxR-type regulators that we observed in this study. We hypothesize that successful HapR inhibitors will require a more targeted approach.

Our modelling experiments successfully predicted the efficacy of the 10A, 10B (PTSP), and 10C molecules, which are the most potent LuxR inhibitors identified thus far. However, it is also clear that the modelling could not reliably predict every critical amino acid contact. Although we identified two critical residues in SmcR, F75 and C170, we hypothesize that more than one critical contact exists because these substitutions did not render HapR sensitive to PTSP. Although the structure of PTSP is similar to that of Qstatin, we were able to show that PTSP is >10X more inhibitory in our assays. We hypothesize that this molecule would make a better initial candidate for future experiments aimed at developing these molecules as therapeutics. In addition to our *in vivo* studies, our *in silico* studies allowed us to model binding affinities of compounds to accurately predict which compounds are best inhibitors of LuxR proteins. This tool will be very useful for future drug development as more compound classes are examined. While modeling has shown to be very effective in our experiments thus far, we recognize that a structure of PTSP bound to SmcR would provide the most information to further optimize *Vibrio* infection therapeutics for PTSP and derivatives.

## Acknowledgements

We thank Stanna Dorn and the Indiana University students in Organic Chemistry C344 laboratory classes for synthesis of molecules in the thiophenesulfonamide panel. We thank the Indiana University School of Medicine Chemical Genomics Core Facility for assistance with chemical screens and analyses. We thank Dr. Irene Newton for assistance with data analysis.

## Author Contributions

JVK and LCB designed the experiments, JVK, JDN, PS, JC, ES, MEM, and LCB performed experiments, JVK, JDN, RH, JC, ES, MEM, and LCB analyzed experimental results, and JVK, JDN, and LCB wrote the manuscript.

## Methods

### Bacterial strains and media

All strains used in this study are listed in Table S3. *E. coli* strains DH10B and S17-1λpir were used for cloning, and BL21(DE3) was used for overexpression of LuxR and SmcR proteins. All *E. coli* strains, *V. cholerae* strains, and derivatives were grown in Lysogeny Broth (LB) at 30°C shaking at 275 RPM in LB media with the appropriate antibiotic. *V. campbellii, V. parahaemolyticus, V. coralliilyticus*, and *V. vulnificus* strains and derivatives were grown shaking at 275 RPM at 30°C in Luria Marine (LM) medium (LB with 2% NaCl) with appropriate antibiotics. Antibiotics were used at the following concentrations: kanamycin 50 μg/mL or 250 μg/mL (*E. coli* or *Vibrio*, respectively), chloramphenicol 10 μg/mL, ampicillin 100 μg/mL, gentamicin 100 μg/mL, and tetracycline 10 μg/mL.

### Molecular methods

All PCR reactions were performed using Phusion HF polymerase (NEB). T4 polynucleotide kinase (T4 PNK) used in EMSAs and all other enzymes were purchased from NEB and used according to manufacturer’s instructions. Site-directed mutagenesis for construction of plasmids expressing mutant proteins was carried out using the Agilent QuikChange II XL Site-Directed Mutagenesis Kit. All oligonucleotides were purchased from Integrated DNA Technologies (IDT), and those used in this study are listed (Table S4). All plasmid constructs (Table S5) were confirmed by DNA sequencing (Eurofins). Cloning details for plasmids are available upon request.

### Compound synthesis and purchase

Synthesis and characterization data for 1B-C, 3A-C, 8A-C, 10A-C, P007 H4, P2065 E16, and P2065 E16-CF3 are provided in the supplementary methods. Compounds P0053 O05 and P1117 F20 were purchased from Lab Network, P1120 D05 and P0074 N08 were purchased from EnamineStore, and P0053 I18 was purchased from ChemDiv.

### E. coli *bioassay*

The dual promoter fluorescence reporter assays were performed using *E. coli* strain DH10B containing two plasmids: 1) plasmid pJV064 containing the P_*luxc*_ fused to GFP and P_*05222*_ fused to mCherry to assess LuxR transcriptional regulation^1^, and 2) plasmid pKM699 expressing *V. campbellii luxR* under control of its native promoter or empty vector pLAFR2^2^. Overnight *E. coli* cultures containing either pKM699 or pLAFR2 and the pJV064 reporter were diluted 1:100 into LB with chloramphenicol and tetracycline and aliquoted into black-welled, clear-bottomed 96-well plates (150 μl final volume). Compounds were resuspended in DMSO and added to *E. coli* cultures at varying concentrations, or DMSO was added as a negative control at equal volumes. 96-well plates were covered in microporous sealing tape and grown for 16 hours shaking at 275 RPM at 30°C. The OD_600_ and fluorescence (both GFP and mCherry) were measured on a BioTek Cytation plate reader.

### Protein purification and electrophoretic mobility shift assays (EMSAs)

SmcR and LuxR were purified as described previously^3^. EMSAs were conducted as described previously^3,4^ using oligonucleotides corresponding to the *luxC* and *vvpE* promoter sequences (Table S4).

### *Assaying compounds in* Vibrio *cultures*

*Vibrio* strains were inoculated in 5 ml LM (or LB for *V. cholerae*) overnight at 30°C shaking at 275 RPM with kanamycin (100 μg/ml) or gentamicin (15 μg/ml) to select for the *PluxC-gfp* reporter plasmids pCS19 or pCS42, respectively. Cultures were back-diluted 1:1,000 in LB or LM with antibiotics, and the cell mixture was aliquoted into black-welled, clear-bottomed 96-well plates. Compounds were titrated into the wells (4-fold dilution series; final volume of 150 μl). DMSO was added as a negative control at equal volumes into control reactions. 96-well plates were covered in microporous sealing tape and grown for 16 hours shaking at 275 RPM at 30°C. The OD_600_ and GFP fluorescence or bioluminescence were measured on a BioTek Cytation plate reader.

### Protease assays

*Vibrio* strains were inoculated in 5 ml LM (or LB for *V. cholerae*) overnight at 30°C shaking at 275 RPM. Cultures were back-diluted 1:1,000 in LB or LM, and the cell mixture was aliquoted into black-welled, clear-bottomed 96-well plates. Compounds were either added into the wells to a specific final concentration or a titration series was performed (4-fold dilution series; final volume of 150 μl; 3 technical replicates per sample). DMSO was added as a negative control at equal volumes into control reactions. 96-well plates were covered in microporous sealing tape and grown for 16 hours shaking at 275 RPM at 30°C. After incubation, the OD_600_ was measured on a BioTek Cytation plate reader. The cultures were pelleted in the 96-well plate by centrifuging at 3700 RPM for 5 min at room temperature. 20 μl of the supernatant was transferred to a new clear 96-well plate. 80 μl of 1% azocasein (dissolved in dH2O) was added to the supernatants and incubated at 37°C for 30 min. 120 μl of 10% trichloracetic acid was added to the reaction, and the plate was incubated on ice for 30 min, then centrifuged at 3700 RPM for 5 min at room temperature. 80 μl of the protease reaction was transferred to a new clear 96-well plate, and 20 μl of 1.8N NaOH was added. The OD_420_ was measured on a BioTek Cytation plate reader. Protease activity was calculated by dividing OD_420_ by OD_600_. Each assay was performed in biological triplicates.

### Autodock Vina modeling and analyses

All docking experiments were performed using Autodock Vina^5^. Xray crystal structures of apo SmcR (PDB ID 3KZ9) and HapR (PDB ID 2PBX) were used for all simulations^6,7^. Structures were prepared for docking using AutoDockTools-1.5.6 for addition of hydrogen atoms and assignment of partial charge^8^. Ligand structures were similarly prepared to include using AutoDockTools which was additionally used to define torsional degrees of freedom. An approximately 14 x 14 x 14 Å box was defined surrounding the residues previously reported to form the Qstatin binding site (box size varied slightly with protein)^9^. Pymol was used visualize the lowest energy solutions. The predicted binding affinities from AutoDock Vina modeling were analyzed compared to the observed IC_50_ values using K-means clustering in R using 3 clusters of sizes 3, 6, and 4.

### Synthesis of LuxR Inhibitors

#### General Methods

^1^H NMR spectra were recorded at room temperature on a Varian I400 (400 MHz) or Varian VXR400 (400 MHz) spectrometer. Chemical shifts are reported in ppm from tetramethylsilane with the residual solvent resonance as the internal standard (CHCl3: δ 7.26 ppm). Data are reported as follows: chemical shift, multiplicity (s = singlet, d = doublet, t = triplet, q = quartet, br = broad, m = multiplet), coupling constants (Hz), and integration. ^13^C NMR spectra were recorded on a Varian I400 (100 MHz) or Varian VXR400 (100 MHz) spectrometer with complete proton decoupling. Chemical shifts are reported in ppm from tetramethylsilane with the solvent resonance as the internal standard (CDCl_3_: δ 77.16 ppm). High Resolution Mass Spectrometry (HRMS) analysis was obtained using Electron Impact Ionization (EI), Chemical Ionization (CI), Atmospheric Pressure Chemical Ionization (APCI) or Electrospray Ionization (ESI) and reported as m/z (relative intensity). ESI was acquired using a Waters/Micromass LCT Classic (ESI-TOF). Dichloromethane (DCM) and tetrahydrofuran (THF) were purified under a positive pressure of dry argon by passage through two columns of activated alumina. Triethylamine (Et3N) and diisopropylethylamine (DIPEA) were distilled over CaH2. All other reagents and solvents were used without purification. All work-up and purification procedures were carried out with reagent grade solvents (purchased from Sigma-Aldrich) in air. Thin-layer chromatography (TLC) was performed on Merck Silica Gel 60 F254 glass plates and visualized with UV and/or standard potassium permanganate, phosphomolybdic acid staining techniques. Standard column chromatography techniques using ZEOprep 60/40-63 μm silica gel was used for purification.

#### General procedure for synthesis of inhibitors 1, 3, 8, 10:^10^

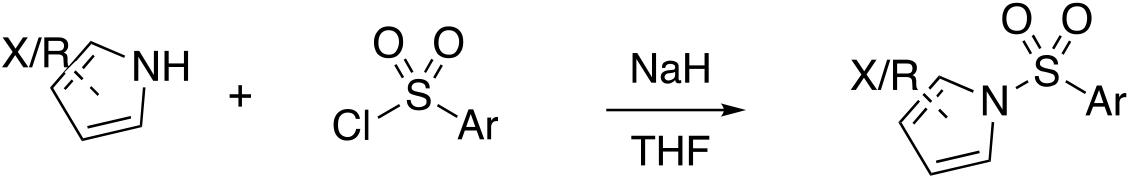

To a solution of amine (6 mmol) in tetrahydrofuran (15 mL) was added sodium hydride (60% in oil, 320 mg, 8 mmol) at room temperature, and the mixture was allowed to stir for 10 min. A solution of sulfonyl chloride (4 mmol) in tetrahydrofuran (5 mL) was added, and the mixture was allowed to stir for an additional 30 min. The reaction mixture was diluted with 20 mL water, and extracted with ethyl acetate (3 x 20 mL). The extract was washed with 20 mL saturated NaCl (brine), dried over anhydrous magnesium sulfate, and concentrated under reduced pressure.

The resulting crude product mixture was purified via SiO_2_ column chromatography in 10:1 Hexanes:EtOAc to give the desired product.

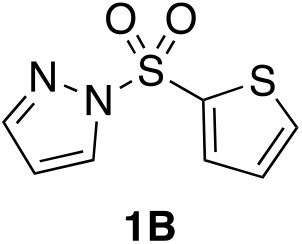

1-(thiophen-2-ylsulfonyl)-1*H*-pyrazole (**1B**): **^1^H NMR (400 MHz, CDCl_3_):** δ 8.08 (d, *J* = 2.7 Hz, 1H), 7.82 (dd, *J* = 3.9, 1.4 Hz, 1H), 7.74 (d, *J* = 1.6 Hz, 1H), 7.71 (dd, *J* = 5.0, 1.4 Hz, 1H), 7.09 (dd, *J* = 5.0, 3.9 Hz, 1H), 6.39 (dd, *J* = 2.8, 1.6 Hz, 1H). **^13^C NMR (101 MHz, CDCl_3_):** δ 145.44, 136.73, 135.51, 135.49, 131.08, 127.87, 109.07. **HRMS (ESI):** Calculated for C_7_H_6_O_2_N_2_NaS_2_ [M+Na^+^]: 236.9763. Found: 236.9763.

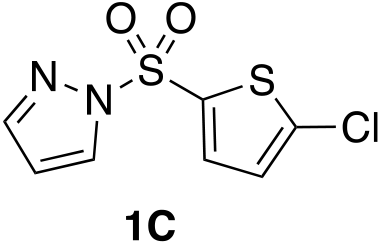

1-((5-chlorothiophen-2-yl)sulfonyl)-1*H*-pyrazole (**1C**): **^1^H NMR (400 MHz, CDCl_3_):** δ 8.06 (d, *J* = 2.8 Hz, 1H), 7.77 (d, *J* = 1.6 Hz, 1H), 7.64 (d, *J* = 4.2 Hz, 1H), 6.94 (d, *J* = 4.2 Hz, 1H), 6.42 (dd, *J* = 2.8, 1.6 Hz, 1H). **^13^C NMR (101 MHz, CDCl_3_):** (101 MHz, cdcl_3_) δ 145.69, 141.32, 134.99, 134.33, 131.10, 127.22, 109.27. **HRMS (APCI):** Calculated for C_7_H_6_ClN_2_O_2_S_2_ [M+H^+^]: 248.9554. Found: 248.9554.

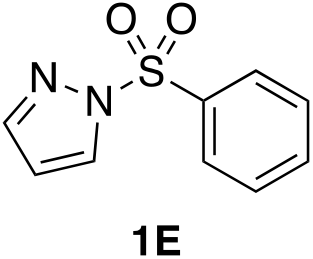

1-(phenylsulfonyl)-1*H*-pyrazole (**1E**): **^1^H NMR (400 MHz, CDCl_3_):** δ 8.10 (d, *J* = 2.8 Hz, 1H), 8.03 - 7.95 (m, 2H), 7.71 (d, *J* = 1.6 Hz, 1H), 7.66 - 7.57 (m, 1H), 7.56 - 7.46 (m, 2H), 6.38 (dd, *J* = 2.8, 1.6 Hz, 1H). **^13^C NMR (101 MHz, CDCl_3_):** δ 145.37, 137.05, 134.55, 131.24, 129.39, 128.03, 108.91. **HRMS (ESI):** Calculated for C_9_HO_2_N_2_NaS [M+Na^+^]: 231.0199. Found: 231.0199.

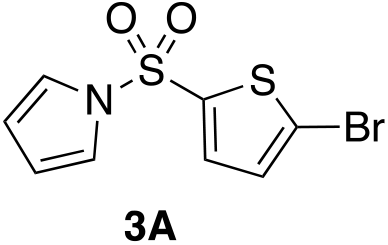

1-((5-bromothiophen-2-yl)sulfonyl)-1*H*-pyrrole (**3A**): **^1^H NMR (400 MHz, CDCl_3_):** δ 7.40 (d, *J* = 4.1 Hz, 1H), 7.13 (t, *J* = 2.3 Hz, 2H), 7.02 (d, *J* = 4.1 Hz, 1H), 6.32 (t, *J* = 2.3 Hz, 2H). **^13^C NMR(101 MHz, CDCl_3_):** δ 139.87, 133.46, 130.52, 122.26, 120.72, 114.31. **HRMS (APCI):** Calculated for C_8_H_7_O_2_NBrS_2_ [M+H^+^]: 291.9096. Found: 291.9098

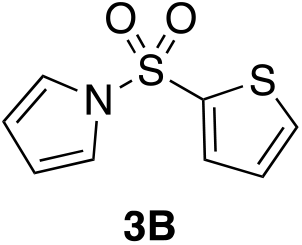

1-(thiophen-2-ylsulfonyl)-1*H*-pyrrole (**3B**): **^1^H NMR (400 MHz, CDCl_3_):** δ 7.63 (ddd, *J* = 13.5, 4.4, 1.4 Hz, 2H), 7.17 (t, *J* = 2.3 Hz, 2H), 7.04 (dd, *J* = 5.1, 3.8 Hz, 1H), 6.30 (t, *J* = 2.3 Hz, 2H). **^13^C NMR (101 MHz, CDCl_3_):** δ 139.39, 133.80, 133.37, 127.54, 120.74, 113.90. **HRMS (EI):** Calculated for C_8_H_7_NO_2_S_2_ [M^+^]: 212.9913. Found: 212.9916.

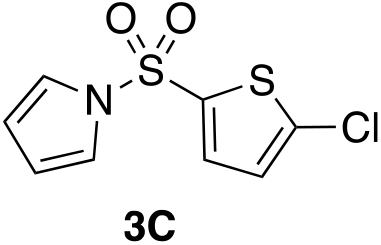

1-((5-chlorothiophen-2-yl)sulfonyl)-1*H*-pyrrole (**3C**): **^1^H NMR (400 MHz, CDCl_3_):** δ 7.44 (d, *J* = 4.1 Hz, 1H), 7.14 (t, *J* = 2.3 Hz, 2H), 6.87 (d, *J* = 4.1 Hz, 1H), 6.32 (t, *J* = 2.3 Hz, 2H). **^13^C NMR (101 MHz, CDCl_3_):** δ 139.51, 136.99, 132.88, 126.99, 120.73, 114.34. **HRMS (EI):** Calculated for C_8_H_6_O_2_NClS_2_ [M^+^]: 246.9528. Found: 246.9533.

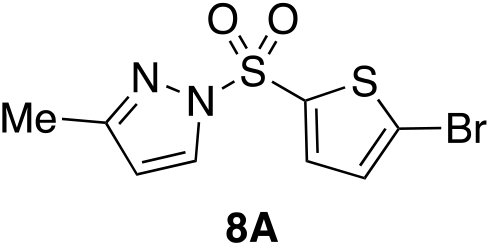

1-((5-bromothiophen-2-yl)sulfonyl)-3-methyl-1*H*-pyrazole (**8A**): **^1^H NMR (400 MHz, CDCl_3_):** δ 7.92 (d, *J* = 2.7 Hz, 1H), 7.55 (d, *J* = 4.1 Hz, 1H), 7.05 (d, *J* = 4.1 Hz, 1H), 6.21 (d, *J* = 2.7 Hz, 1H), 2.28 (s, 3H). **^13^C NMR (101 MHz, CDCl_3_):** δ 156.08, 137.68, 135.16, 132.02, 130.69, 123.51, 110.25, 14.01. **HRMS (ESI):** Calculated for C_8_H_7_O_2_N_2_BrNaS_2_ [M+Na^+^]: 330.9003. Found: 330.9004.

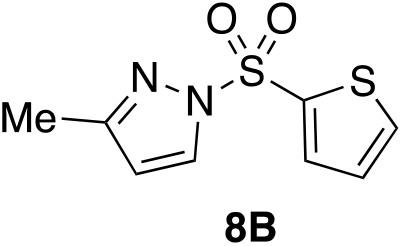

3-methyl-1-(thiophen-2-ylsulfonyl)-1*H*-pyrazole (**8B**): **^1^H NMR (400 MHz, CDCl_3_):** δ 7.96 (d, *J* = 2.7 Hz, 1H), 7.80 (dd, *J* = 3.9, 1.4 Hz, 1H), 7.68 (dd, *J* = 5.0, 1.4 Hz, 1H), 7.08 (dd, *J* = 5.0, 3.8 Hz, 1H), 6.20 (d, *J* = 2.7 Hz, 1H), 2.27 (s, 3H). **^13^C NMR (101 MHz, CDCl_3_):** δ 155.70, 137.22, 135.06, 134.93, 131.99, 127.71, 109.96, 13.98. **HRMS (ESI):** Calculated for C_8_H_8_N_2_O_2_S_2_Na [M+Na^+^]: 250.9919. Found: 250.9920.

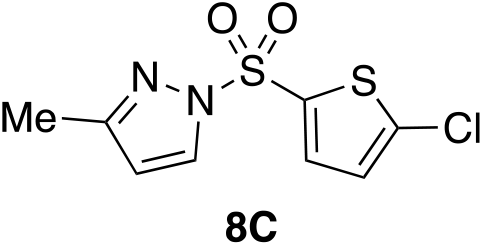

1-((5-chlorothiophen-2-yl)sulfonyl)-3-methyl-1*H*-pyrazole (**8C**): **^1^H NMR (400 MHz, CDCl_3_):** δ 7.92 (d, *J* = 2.7 Hz, 1H), 7.59 (d, *J* = 4.2 Hz, 1H), 6.91 (d, *J* = 4.1 Hz, 1H), 6.21 (d, *J* = 2.8 Hz, 1H), 2.28 (s, 3H). **^13^C NMR (101 MHz, CDCl_3_):** δ 156.08, 140.71, 134.79, 134.54, 132.02, 127.10, 110.25, 14.00. **HRMS (ESI):** Calculated for C_8_H_7_O_2_N_2_ClNaS_2_ [M+Na^+^]: 283.9530. Found: 284.9531.

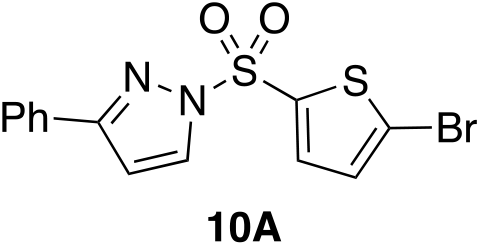

1-((5-bromothiophen-2-yl)sulfonyl)-3-phenyl-1*H*-pyrazole (**10A**): **^1^H NMR (400 MHz, CDCl_3_):** δ 8.08 (d, *J* = 2.8 Hz, 1H), 7.87 - 7.79 (m, 2H), 7.61 (d, *J* = 4.1 Hz, 1H), 7.45 - 7.37 (m, 3H), 7.05 (d, *J* = 4.1 Hz, 1H), 6.73 (d, *J* = 2.8 Hz, 1H). **^13^C NMR (101 MHz, CDCl_3_):** δ 157.47, 137.37, 135.42, 132.53, 131.12, 130.73, 129.48, 128.74, 126.49, 123.94, 107.10. **HRMS (ESI):** Calculated for C_13_H_9_BrN_2_O_2_S_2_Na [M+Na^+^]: 390.9181. Found: 390.9182

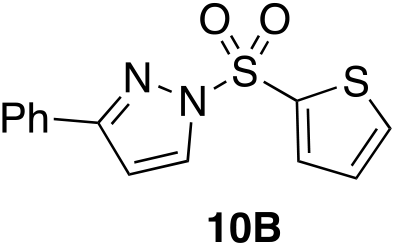

3-phenyl-1-(thiophen-2-ylsulfonyl)-1*H*-pyrazole (**10B**): **^1^H NMR (400 MHz, CDCl_3_):** δ 8.11 (d, *J* = 2.8 Hz, 1H), 7.88 - 7.79 (m, 3H), 7.68 (dd, *J* = 5.0, 1.4 Hz, 1H), 7.44 - 7.31 (m, 3H), 7.07 (dd, *J* = 5.0, 3.9 Hz, 1H), 6.71 (d, *J* = 2.8 Hz, 1H). **^13^C NMR (101 MHz, CDCl_3_):** δ 157.17, 136.84, 135.40, 135.36, 132.55, 131.28, 129.36, 128.72, 127.79, 126.47, 106.90. **HRMS (ESI):** Calculated for C_13_H_10_O_2_N_2_NaS_2_ [M+Na^+^]: 313.0076. Found: 313.0078.

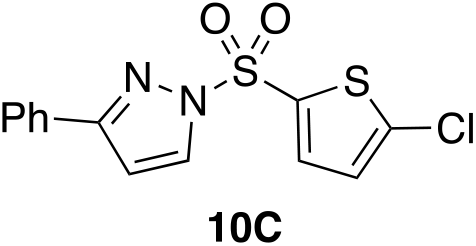

1-((5-chlorothiophen-2-yl)sulfonyl)-3-phenyl-1*H*-pyrazole (**10C**): **^1^H NMR (400 MHz, CDCl_3_):** δ 8.08 (d, *J* = 2.8 Hz, 1H), 7.87 - 7.79 (m, 2H), 7.65 (d, *J* = 4.2 Hz, 1H), 7.41 (s, 1H), 7.43 – 7.33 (m, 2H), 6.92 (d, *J* = 4.1 Hz, 1H), 6.73 (d, *J* = 2.8 Hz, 1H). **^13^C NMR (101 MHz, CDCl_3_):** δ 157.47, 141.13, 134.79, 134.50, 132.51, 131.12, 129.47, 128.74, 127.12, 126.48, 107.08. **HRMS (ESI):** Calculated for C_13_H_9_O_2_N_2_ClNaS_2_ [M+Na^+^]: 346.9686. Found: 346.9688.

### Synthesis of P007H4^11^

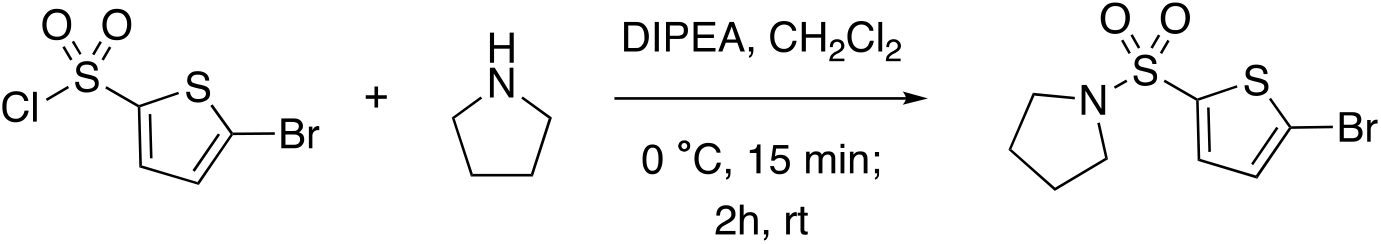

The sulfonyl chloride (1.05 g, 4 mmol) was dissolved in dichloromethane (1.0 mL) and the resulting solution was added dropwise to a round-bottom flask containing a stirred solution of pyrrolidine (657 μL, 8 mmol) and diisopropylethyllamine (1.5 mL, 8 mmol) in dichloromethane (10 mL) at 0 °C. The resulting reaction mixture was allowed to stir at 0 °C for 15 min and then allowed to warm to room temperature. After 2 h, the resulting solution was washed with 10 mL each of saturated sodium bicarbonate, water, 1 N HCl and brine. The organic layer was dried with magnesium sulfate, and solvent was removed under reduced pressure to give 1-((5-bromothiophen-2-yl)sulfonyl)pyrrolidine (**P007H4**). **^1^H NMR (400 MHz, CDCl_3_):** δ 7.30 (d, *J* = 4.0 Hz, 1H), 7.08 (d, *J* = 4.0 Hz, 1H), 3.29 - 3.22 (m, 4H), 1.85 - 1.74 (m, 4H). **^13^C NMR (101 MHz, CDCl_3_):** δ 137.98, 132.05, 130.47, 119.30, 48.20, 25.34. **HRMS (ESI):** Calculated for C_9_H_10_BrNO_2_S_2_Na [M+Na^+^]: 317.9229. Found: 317.9229.

### Synthesis of P2065E16

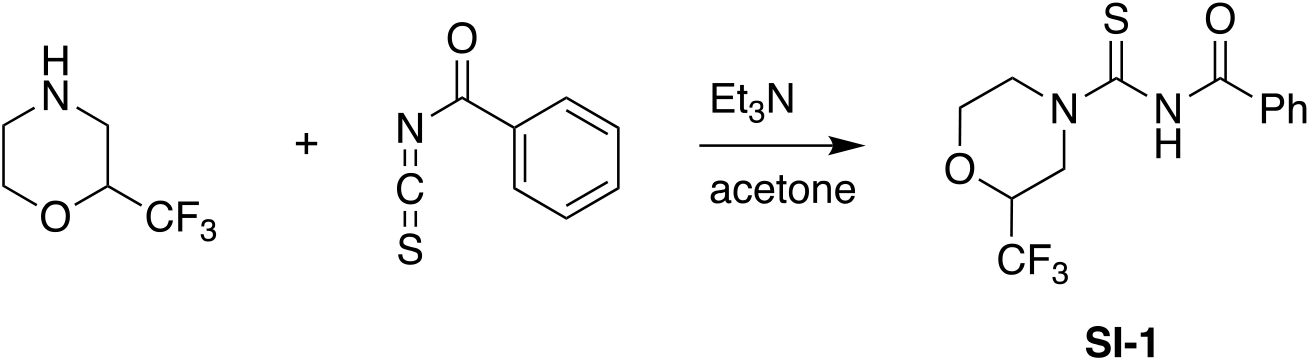

To a flask purged and backfilled under nitrogen was added a solution of 2-trifluoromethylmorpholine (100 mg, 0.522 mmol) in 0.5 mL dry acetone. Triethylamine (0.109 mL, 0.783 mmol) was added via syringe and the resulting solution allowed to stir at room temperature for 30 minutes. The solution was cooled to 0°C, and benzoyl isothiocyanate (0.702 mL, 0.522 mmol) was added dropwise. The resulting solution was allowed to stir for 30 min at 0°C and was then quenched with 1 mL water. The crude product was extracted with ethyl acetate (2 x 30 mL). The organic layers were combined, dried over anhydrous sodium sulfate, and concentrated under reduced pressure. The resulting crude product mixture was purified via SiO_2_ column chromatography in 5:1 Hexanes:EtOAc to give *N*-(2-(trifluoromethyl)morpholine-4-carbonothioyl)benzamide (**SI-1**) in 62% yield^12^. **^1^H NMR (400 MHz, CDCl_3_):** δ 7.93 - 7.75 (m, 2H), 7.69 - 7.55 (m, 1H), 7.50 (t, *J* = 7.7 Hz, 2H), 5.10 (d, *J* = 94.4 Hz, 2H), 4.11 (s, 3H), 3.86 (s, 1H), 3.58 - 3.42 (m, 1H), 3.37 (dd, *J* = 13.4, 10.6 Hz, 1H).

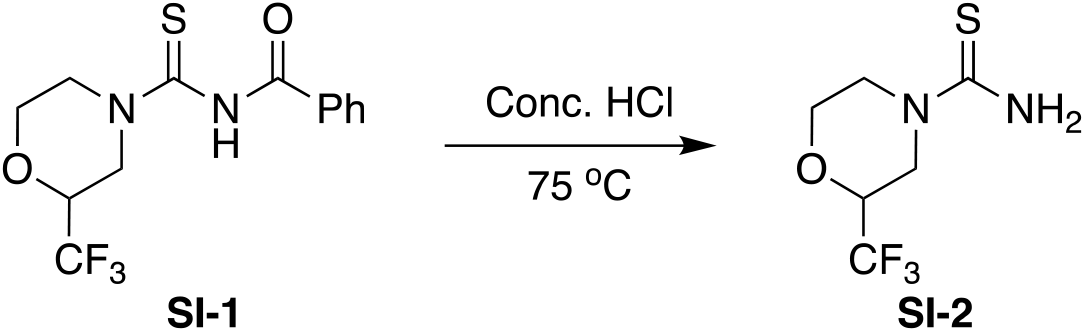

Concentrated HCl (2 mL) was added to **SI-1** (102.6 mg, 0.322 mmol). The resulting solution was allowed to stir for 1.5 hours at 75°C, then cooled to 0°C. Water (10 mL) was added, followed by 50% sodium hydroxide (5 mL). The aqueous phase was extracted with 1:1 ethyl acetate: petroleum ether (3 x 30mL). The organic layers were combined, washed with water (10 mL), dried over anhydrous sodium sulfate, and concentrated under reduced pressure. The resulting crude product mixture was purified via SiO_2_ column chromatography in 1:1 Hexanes:EtOAc to give 2-(trifluoromethyl)morpholine-4-carbothioamide (**SI-2**) in 51% yield^13^. **^1^H NMR (400 MHz, CDCl_3_):** δ 6.33 (s, 2H), 4.75 (d, *J* = 13.3 Hz, 1H), 4.30 (d, *J* = 13.4 Hz, 1H), 3.92 (dqd, *J* = 12.1, 6.0, 3.0 Hz, 1H), 3.64 (td, *J* = 11.6, 3.0 Hz, 1H), 3.25 (ddd, *J* = 13.5, 11.3, 3.6 Hz, 1H), 3.18 (dd, *J* = 13.4, 10.6 Hz, 1H).

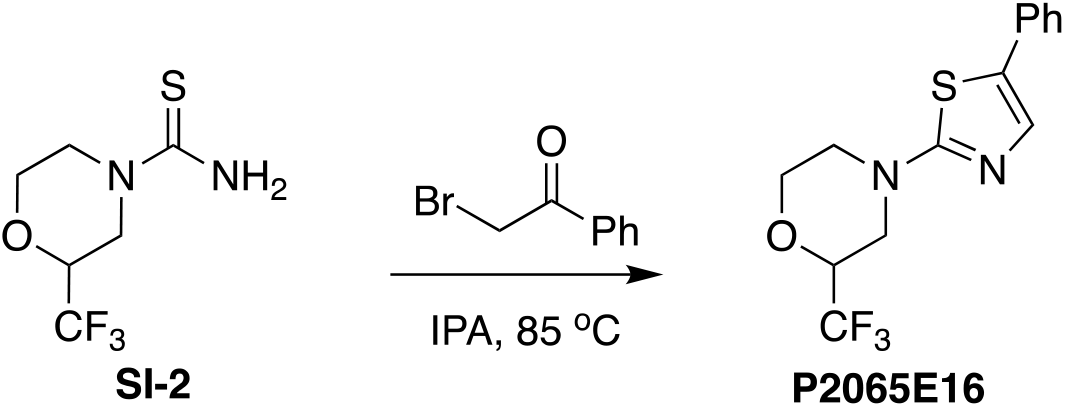

**SI-2** (35 mg, 0.164 mmol) and 2-bromoacetophenone (33 mg, 0.164 mmol) were dissolved in isopropanol (5 mL). The resulting solution was heated with stirring for four hours at 85°C. The reaction mixture was cooled to room temperature and concentrated under reduced pressure. The resulting crude reaction mixture was purified via SiO_2_ column chromatography (hexanes/EtOAc) to give 4-(5-phenylthiazol-2-yl)-2-(trifluoromethyl)morpholine (**P2065E16**) in 24% yield^14^. **^1^H NMR (400 MHz, CDCl_3_):** δ 7.76 (d, *J* = 7.4 Hz, 2H), 7.32 (t, *J* = 7.7 Hz, 2H), 7.19 (d, *J* = 2.2 Hz, 1H), 6.79 (s, 1H), 4.10 - 3.91 (m, 3H), 3.75 (t, *J* = 15.1 Hz, 2H), 3.18 (dt, *J* = 38.2, 11.9 Hz, 2H). **HRMS (APCI):** Calculated for C_14_H_13_F_3_N_2_OS [M ^+^]: 314.0701. Found: 314.0702.

### Synthesis of P2065E16-CF_3_

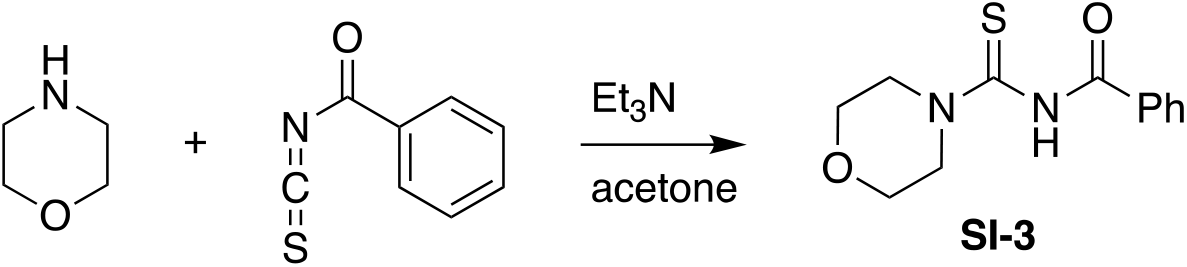

Morpholine (1.0 g, 0.99 mL, 11.5 mmol) was added to dry acetone (10 mL) under nitrogen. Triethylamine (1.74 g, 2.40 mL, 17.2 mmol) was added via syringe and the resulting solution was allowed to stir at room temperature for 30 minutes. The resulting solution was cooled to 0°C, and benzoyl isothiocyanate (1.87 g, 1.54 mL, 11.5 mmol) was added dropwise. The resulting solution was allowed to stir for 30 min at 0°C and was then quenched with water (20 mL). The aqueous layer was extracted with ethyl acetate (2 x 30 mL). The organic layers were combined, dried over anhydrous sodium sulfate, and concentrated under reduced pressure. The resulting crude product mixture was purified via SiO_2_ column chromatography in 3:1 Hexanes:EtOAc to give *N*-(morpholine-4-carbonothioyl)benzamide (**SI-3**) in 57% yield. **^1^H NMR (400 MHz, CDCl_3_):** δ 8.55-8.50 (m, 1H), 7.81 (d, J = 7.6 Hz, 2H), 7.57 (t, J = 7.6 Hz, 1H), 7.46 (t, J = 7.6 Hz, 2H), 4.20 (s, 2H), 3.85 - 3.77 (m, 4H), 3.63 (s, 2H).

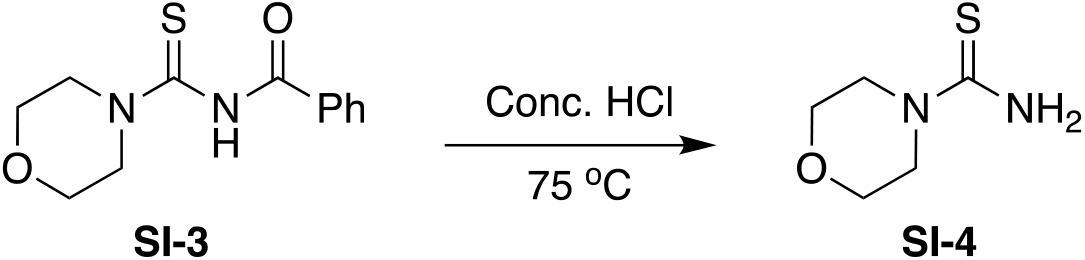

Concentrated HCl (4 mL) was added to **SI-3** (169 mg, 0.675 mmol). The resulting solution was heated with stirring for 1.5 hours at 75°C, then cooled to 0°C. Water (15 mL) was added, followed by 50% sodium hydroxide (10 mL). The aqueous phase was extracted with 1:1 ethyl acetate: petroleum ether (3 x 30 mL). The combined organic layers were dried over anhydrous sodium sulfate and concentrated under reduced pressure to provide morpholine-4-carbothioamide (**SI-4**) in 33% yield (no purification was necessary). **^1^H NMR (400 MHz, CDCl_3_):** δ 7.42 (s, 2H), 3.67 (t, *J* = 4.8 Hz, 4H), 3.52 (t, *J* = 4.8 Hz, 4H).

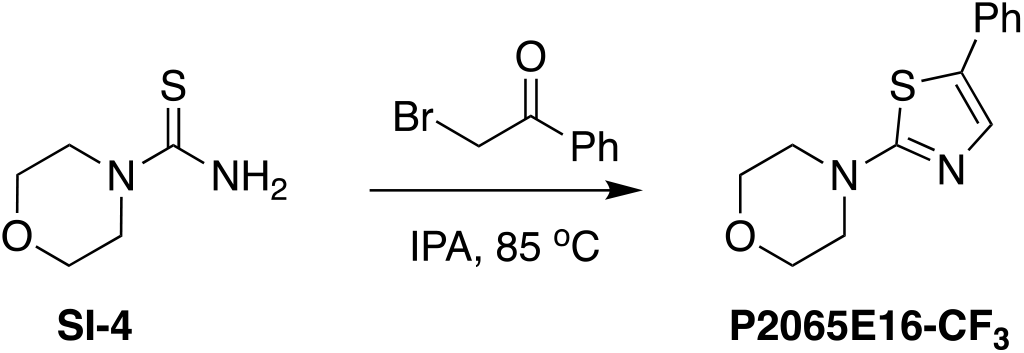

**SI-4** (123 mg, 0.84 mmol) and 2-bromoacetophenone (167 mg, 0.84 mmol) were dissolved in isopropanol (5 mL). The resulting solution was heated with stirring for four hours at 85°C. The reaction was cooled to room temperature and concentrated under reduced pressure to give 4- (5-phenylthiazol-2-yl)morpholine (**P2065E16-CF_3_**) in quantitative yield. δ 7.82 - 7.76 (m, 2H), 7.37 (t, *J* = 7.6 Hz, 2H), 7.31-7.25 (m, 2H), 3.70 (t, *J* = 4.9 Hz, 4H), 3.46 (t, *J* = 4.8 Hz, 4H). **^13^C NMR (101 MHz, DMSO-d_6_):** δ 170.99, 148.82, 133.74, 129.02, 128.49, 126.42, 103.43, 65.71, 48.87. **HRMS (EI):** Calculated for C_13_H_14_N_2_OS [M ^+^]: 246.0827. Found: 246.0839.

