## Supplementary Info for "Thiophenesulfonamides are specific inhibitors of quorum sensing in pathogenic Vibrios"

**Supplementary Information**
Tables S1-S5
Figures S1-S6

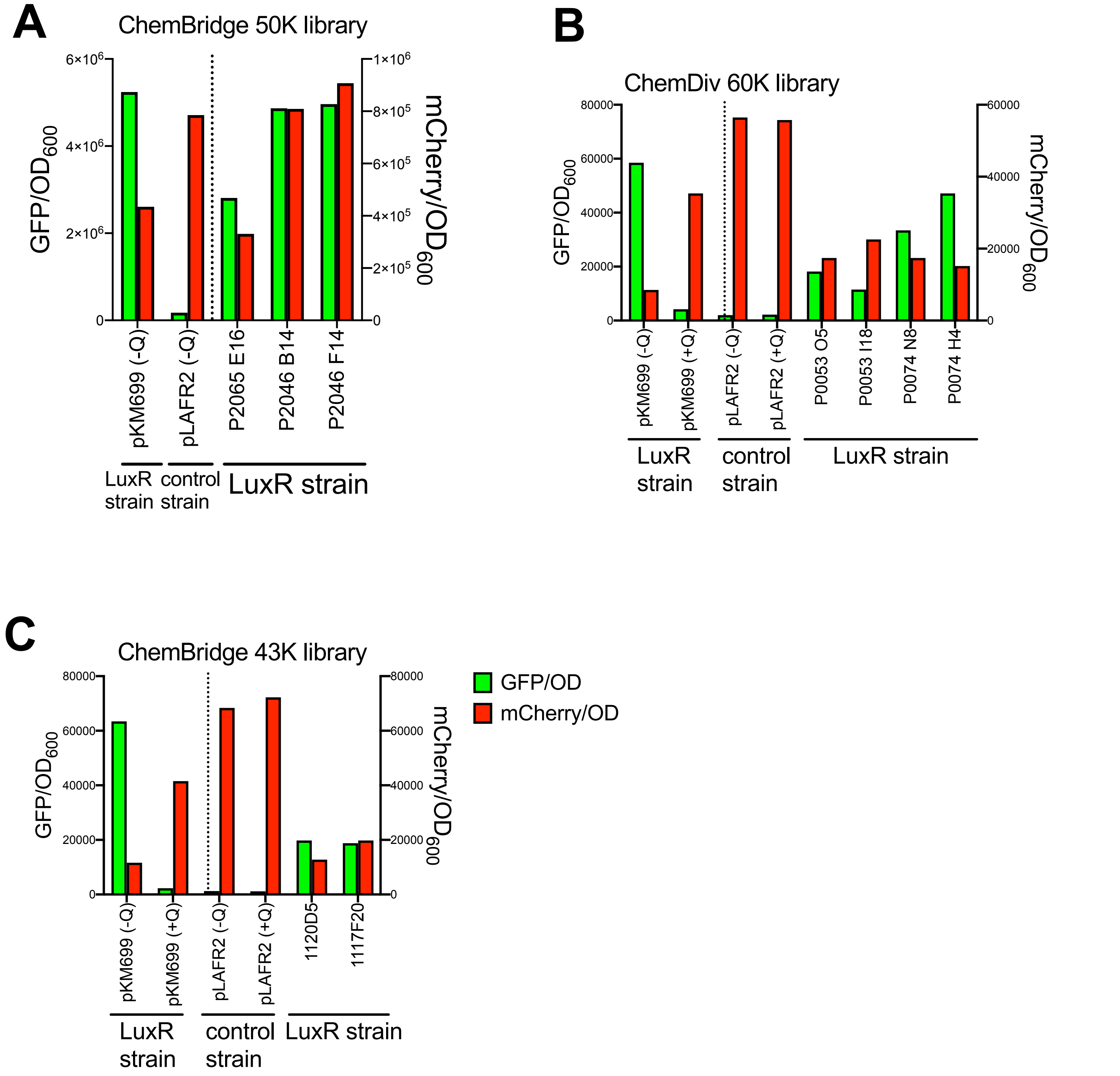


**Supplemental Figure 1.** (A-C) Production of GFP (GFP/$OD_{600}$) or mCherry (mCherry/$OD_{600}$) in the presence of compounds in chemical libraries (final concentration is 12.5 μg/ml) in the *E. coli* bioassay strain containing plasmid pKM699 expressing LuxR (pJV064). Qstatin (Q+) was added at a final concentration of 5 μM (1.47 μg/ml) compared to the solvent DMSO (Q-) as positive and negative controls, respectively. The control strain contains empty vector control pLAFR2 instead of pKM699.

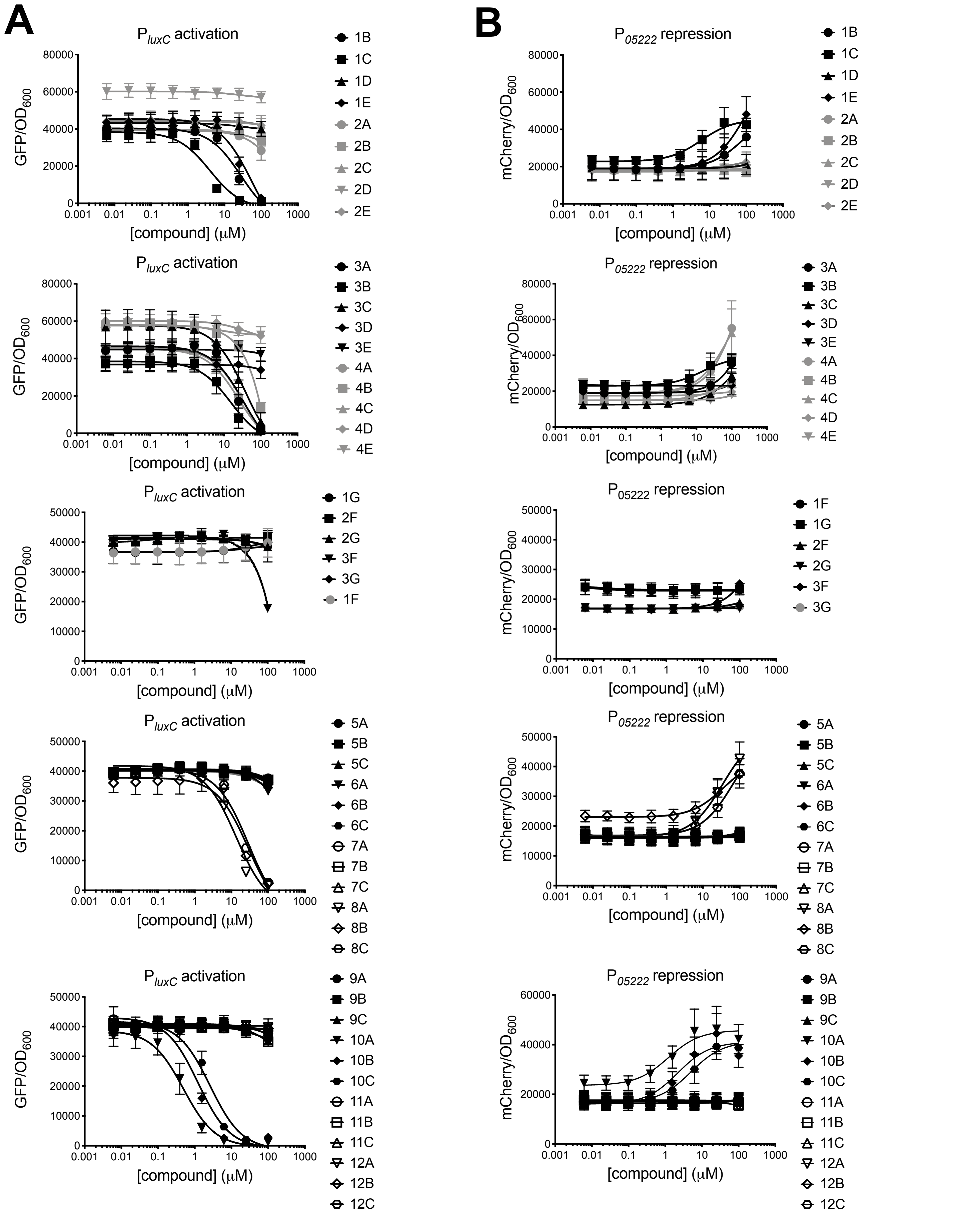


**Supplemental Figure 2.** Production of GFP (panel A; GFP/$OD_{600}$) or mCherry (panel B; mCherry/$OD_{600}$) in the presence of sulfonamide compounds titrated into the *E. coli* bioassay strain (pKM699, pJV064). Data shown represent the mean and standard deviation of at least three biological replicates.

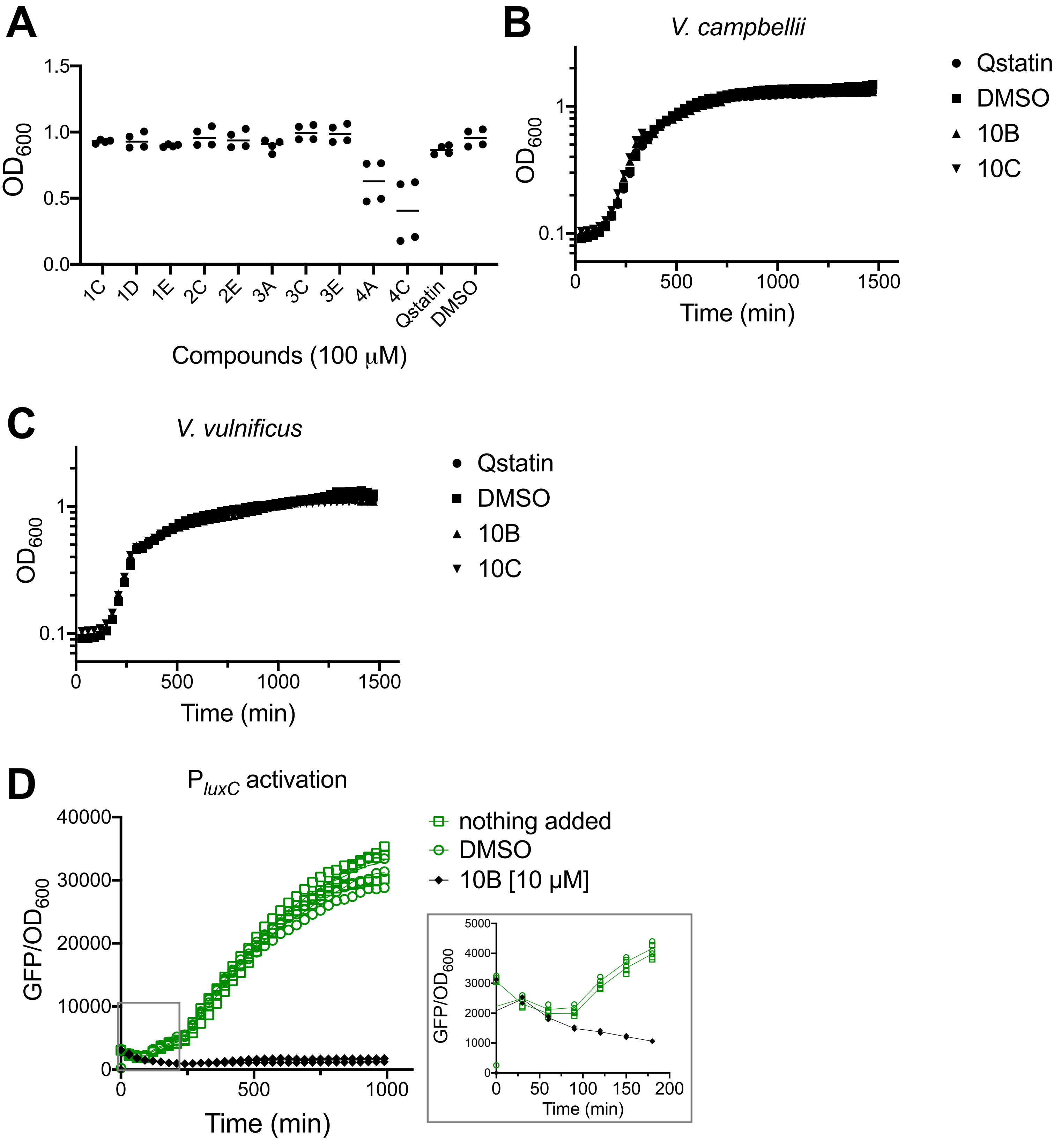


**Supplemental Figure 3.** (A) Growth yield of *E. coli* bioassay strain (pKM699, pJV064) in the presence of compounds at 100 μM or an equal volume of DMSO. (B, C) Growth curves of *V. campbellii* BB120 (B) and *V. vulnificus* ATCC 27562 (C) in the presence of 25 μM 10B, 10C, Qstatin, or an equal volume of DMSO. (D) Production of GFP (GFP/$OD_{600}$) over a timecourse in the presence of 10 μM compound 10B added at time 0 to the *E. coli* bioassay strain (pKM699, pJV064). An equal volume of solvent DMSO was added as a negative control compared to nothing added. Inset is the first 180 minutes of the timecourse.

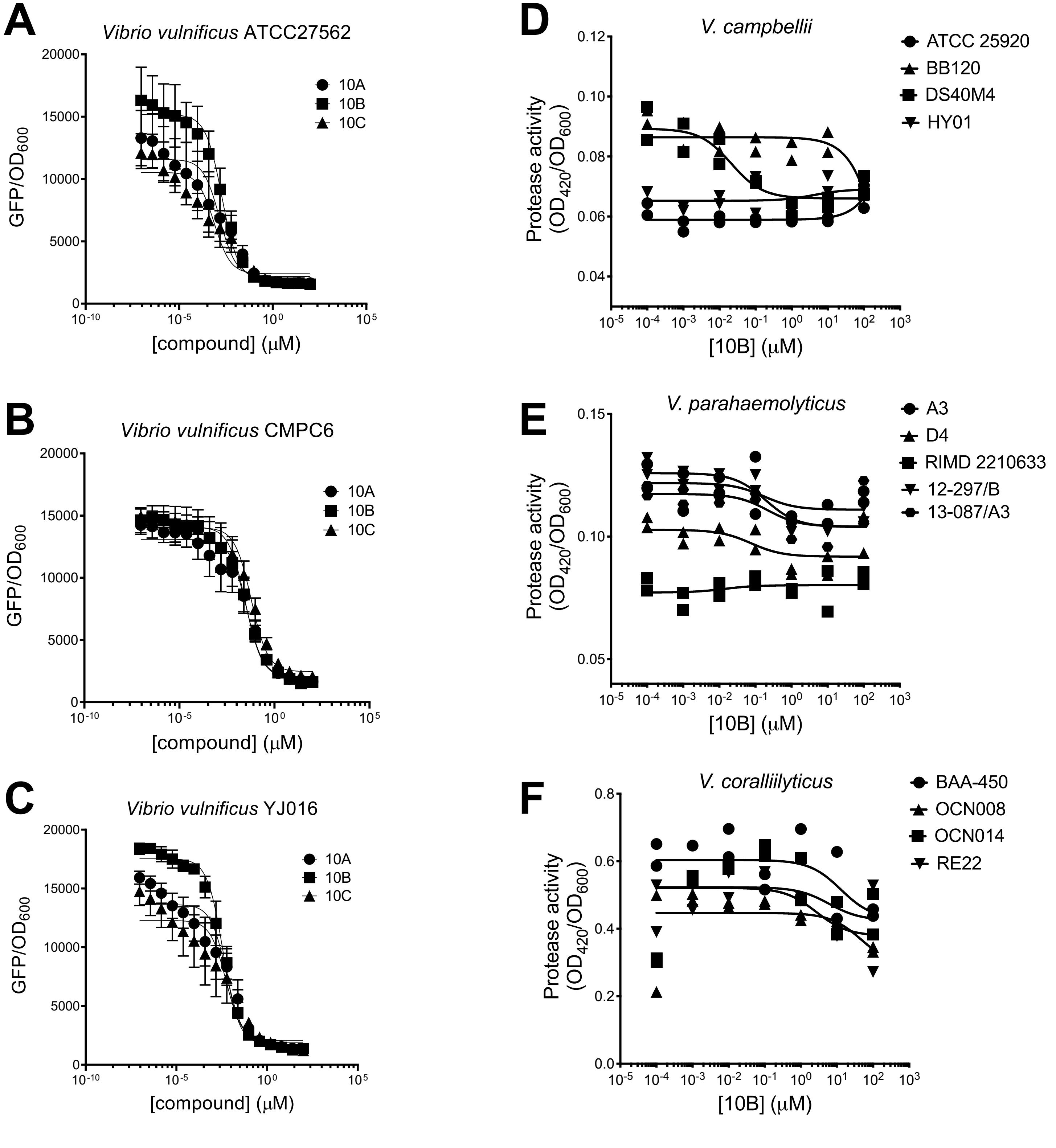


**Supplemental Figure 4**. (A-C) Titration of molecules from the top panel of thiophenesulfonamides in *V. vulnificus* strains. Data shown represent the mean and standard error of measurement of three biological replicates. (D-F) Protease activity (final assay $OD_{420}$/initial culture $OD_{600}$) for *Vibrio* strains in the presence of 10B titrated into the cultures. Two biological replicates are shown for each culture series.

PTSP modelling in SmcR and HapR

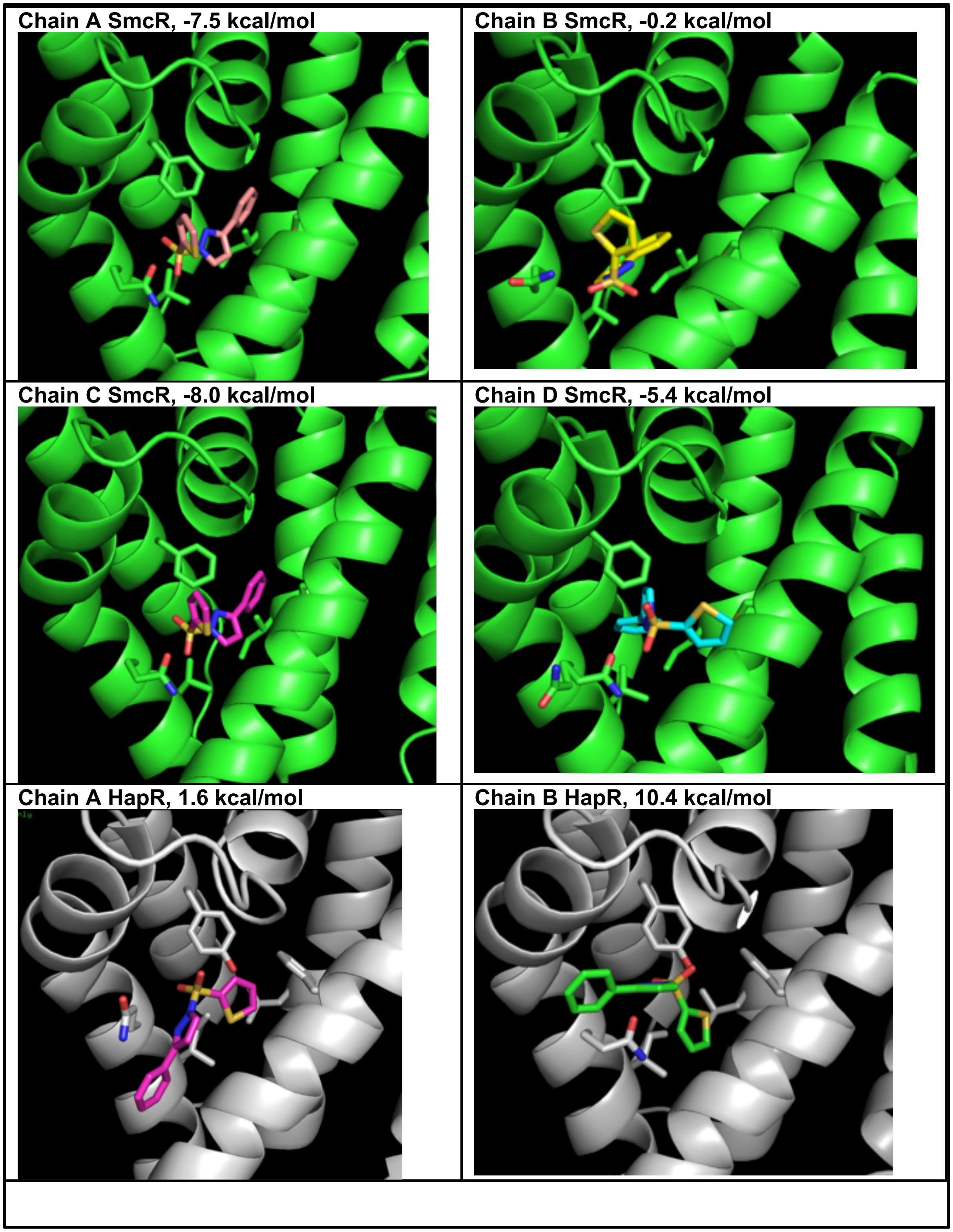


**Supplemental Figure 5**. Autodock Vina modelling of PTSP into the four chains of SmcR (3KZ9) and two chains of HapR (2PBX). The predicted binding energies are shown.

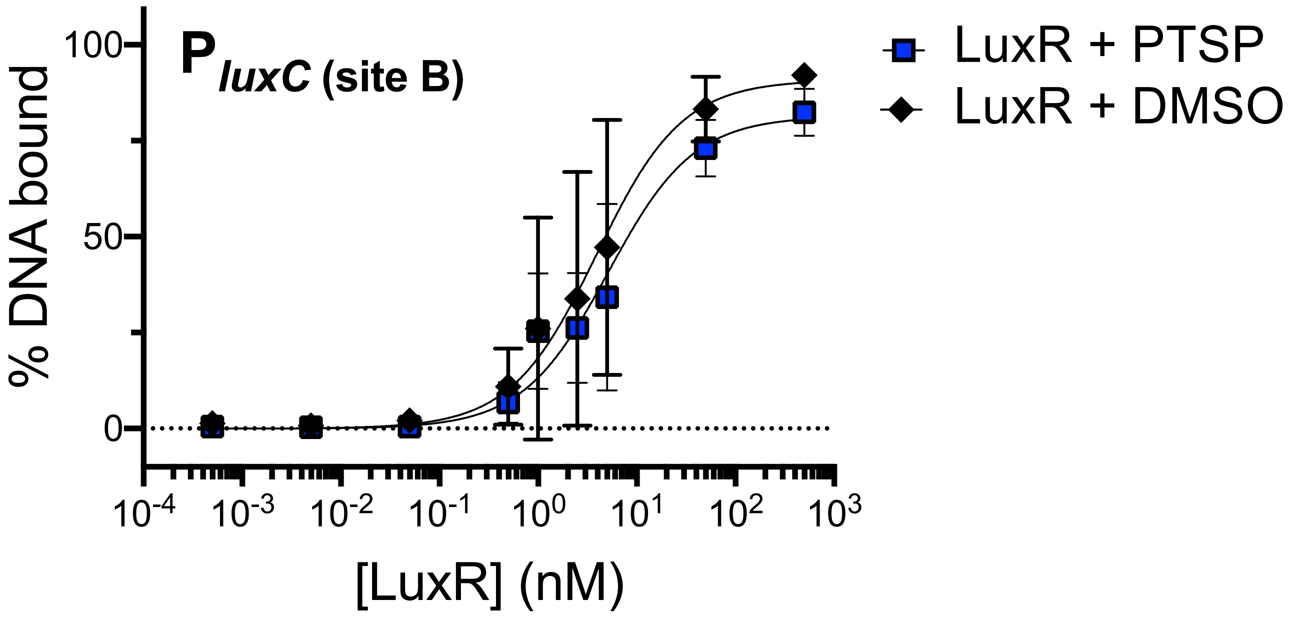


**Supplemental Figure 6**. EMSAs with purified LuxR protein in varying concentrations incubated with radiolabeled dsDNA corresponding to the *V. campbellii luxC* promoter site B. DNA shifts were quantified using ImageJ, and the graphs show the mean and standard deviation for three biological replicates.

**Tables**

**Table S1. Inhibition ($IC_{50}$) of GFP production by LuxR in *E. coli* bioassay.**

| Molecule | $IC_{50}$ (μM) | $IC_{50}$ 95% confidence interval (μM) |
|---|---|---|
| P0053 I18 | 1.9 | 1.2 – 3.0 |
| P0053 O05 | 16.5 | 12.0 – 22.8 |
| P0074 H04 | ND | ND |
| P0074 N08 | 14.6 | 8.1 – 26.9 |
| P1032 E02 | ND | ND |
| P1117 F20 | 1.1 | 0.2 – 4.1 |
| P1120 D05 | 73.7 | 40.7 – 171.2 |
| P2046 F14 | ND | ND |
| P2065 E16 | 5.3 | 2.8 – 9.9 |
| Qstatin | 5.0 | 3.5 – 7.0 |

ND = a curve could not be fit to these data to determine $IC_{50}$.

**Table S2. Inhibition ($IC_{50}$) of the $P_{luxC\text{-}gfp}$ reporter in *Vibrio* strains.**

| $IC_{50}$ (μM)[a] | Qstatin | 1B | 1C | 1E | 3A | 3B | 3C[b] | 8A | 8B | 8C | 10A | 10B | 10C | P0053 I18 |
|---|---|---|---|---|---|---|---|---|---|---|---|---|---|---|
| ***Vcamp*** **BB120** | 4.0 | 52.6 | 3.3 | 51.4 | 15.0 | 18.9 | 8.3 | 3.8 | 32.4 | 6.7 | 0.58 | 0.35 | 0.71 | 2.0 |
| ***Vcamp*** **HY01** | 1.2 | 23.2 | 1.0 | 14.8 | 1.6 | 3.6 | ND | 1.2 | 8.7 | 2.7 | 0.40 | 0.10 | 0.16 | 4.3 |
| ***Vcamp*** **ATCC 25920** | 0.8 | 12.8 | 0.5 | 8.4 | 1.1 | 2.0 | ND | 0.7 | 3.9 | 1.5 | 0.05 | 0.03 | 0.09 | 1.8 |
| ***Vcoral*** **OCN008** | 11.6 | 55.7 | 9.1 | 49.2 | 23.1 | 39.2 | 1.3 | 4.2 | 26.1 | 2.7 | 2.09 | 1.43 | 1.07 | 18.9 |
| ***Vcoral*** **OCN014** | 22.2 | 100.6 | 9.4 | 66.2 | 22.6 | 95.1 | ND | 9.6 | 26.8 | 8.4 | 2.01 | 2.37 | 2.42 | 10.4 |
| ***Vpara*** **RIMD 2210633** | 2.3 | 15.9 | 1.7 | 11.0 | 2.6 | 2.5 | 2.0 | 2.0 | 7.3 | 3.8 | 0.63 | 0.17 | 0.68 | 2.8 |
| ***Vpara*** **D4** | 3.0 | 18.9 | 2.5 | 12.7 | 2.8 | 3.5 | ND | 2.4 | 8.4 | 4.5 | 0.87 | 0.27 | 1.28 | 6.4 |
| ***Vpara*** **12-297/B** | 1.7 | 11.8 | 1.6 | 8.1 | 3.0 | 4.0 | ND | 2.1 | 6.7 | 3.3 | 1.03 | 0.32 | 1.04 | 8.1 |
| ***Vvul*** **ATCC 27562** | 0.5 | 3.4 | 0.4 | 2.7 | 0.7 | 0.9 | 0.3 | 1.9 | 0.7 | 0.0 | 0.00144 | 0.00196 | 0.00075 | 0.7 |
| ***Vvul*** **CMPC6** | 2.1 | 29.6 | 2.3 | 22.0 | 6.1 | 5.9 | ND | 2.1 | 11.5 | 4.6 | 0.03392 | 0.02949 | 0.06046 | 4.9 |
| ***Vvul*** **YJ016** | 0.9 | 6.5 | 0.8 | 4.6 | 1.3 | 1.5 | ND | 0.5 | 3.0 | 1.0 | 0.00584 | 0.00378 | 0.00546 | 3.2 |

a. *Vcamp*, *V. campbellii*; *Vcoral, V. coralliilyticus*; *Vpara, V. parahaemolyticus*; *Vvul, V. vulnificus*
b. ND, not determined.

**Table S3. Bacterial strains used in this study.**

| Name | Genotype/Notes | Reference |
|---|---|---|
| ***E. coli* strains** | | |
| S17-1λ*pir* | Conjugation strain, λ-*pir*, *recA thi pro hsdR*$^-$ *M*$^+$ RP4: 2-Tc, Mu, Tp$^R$, Sm$^R$ | [1] |
| DH10B | str. K-12 F$^-$ Δ(*ara-leu*)*7697*[Δ(*rapA'-cra'* )] Δ(*lac*)*X74*[Δ('*yahH-mhpE*)] duplication (514341-627601) [*nmpC-gltI*] *galK16 galE15* e14$^-$(*icd*$^{WT}$ *mcrA*) φ80d*lacZΔM15 recA1 relA1 endA1* Tn*10.10 nupG rpsL150* (Str$^R$) *rph*$^+$ *spoT1* Δ*(mrr-hsdRMS- mcrBC)* λ$^-$ Missense (*dnaA glmS glyQ lpxK mreC murA*) Nonsense (*chiA gatZ fhuA*? *yigA ygcG*) Frameshift(*flhC mglA fruB*) | Life Technologies |
| BL21(DE3) | *E. coli* str. B F$^-$ *ompT gal dcm lon hsdS*$_B$($r_B^-m_B^-$) λ(DE3 [*lacI lacUV5-T7p07 ind1 sam7 nin5*]) [*malB*$^+$]$_{K\text{-}12}$(λ$^S$) | NEB |
| β3914 | Conjugation strain; Δ*dapA*::(*erm-pir*); Km$^r$, Em$^r$, Tc$^r$ | [2] |
| ***V. campbellii* strains** | | |
| BB120 | Wild-type | [3] |
| KM669 | BB120 Δ*luxR* | [4] |
| HY01 | Wild-type | [5] |
| ATCC 25920 | (NBRC 15631/ CAIM 519) | ATCC |
| ***V. coralliilyticus* strains** | | |
| OCN008 | Wild-type | [6,7] |
| Δ*vcpR* | OCN008 Δ*vcpR* | [8] |
| OCN014 | Wild-type | [9] |
| ***V. cholerae* strains** | | |
| E7946 | wild-type 01 El Tor | [10] |
| SAD793 | E7946 Sm$^R$ Δ*hapR::Spec*$^R$, pMMB67EH-tfox-kanR | [11] |
| ***V. parahaemolyticus* strains** | | |
| RIMD 2210633 | Wild-type | ATCC |
| CAS-V03 | RIMD2210633 Δ*opaR::Spec*$^R$, pMMB67EH-tfoX-kanR | [11] |
| D4 | AHPND isolate | [12] |
| 12-197/B | AHPND isolate | [12] |
| ***V. vulnificus* strains** | | |
| ATCC 27562 | Wild-type | ATCC |
| CAS-vv015 | ATCC 27562 Δ*smcR* | [11] |
| YJ016 | Clinical isolate | [13] |
| CMPC6 | Clinical isolate | [14] |

**Table S4. Oligonucleotides used in this study.**

| Name | Description | Reference |
|---|---|---|
| JCV1295 | GATAACAATTTCACACAGGAAACAGAATTCATGGACTCAATTGCAAAGAGAC | luxR in pMMB, F |
| JCV1296 | GCATGCCTGCAGGTCGACTCTAGAGGATCCTTAGTGATGTTCACGGTTGTAGA | luxR in pMMB, R |
| JCV1297 | GATAACAATTTCACACAGGAAACAGAATTCATGGACGCATCAATCGAAAAAC | hapR n pMMB, F |
| JCV1298 | GCATGCCTGCAGGTCGACTCTAGAGGATCCCTAGTTCTTATAGATACACAGCATATTGA | hapR in pMMB, R |
| JCV1299 | GATAACAATTTCACACAGGAAACAGAATTCATGGACTCAATTGCAAAGAGAC | opaR in pMMB, F |
| JCV1300 | GCATGCCTGCAGGTCGACTCTAGAGGATCCTTAGTGTTCGCGATTGTAGATGC | opaR in pMMB, R |
| JCV1301 | GATAACAATTTCACACAGGAAACAGAATTCATGGATTCTATAGCTAAGAGACCG | vcpR in pMMB, F |
| JCV1302 | GCATGCCTGCAGGTCGACTCTAGAGGATCCCTACTTGTAGATGCAAAGCATATCTAG | vcpR in pMMB, R |
| JCV1305 | AAGTGCTTAATCATGTCGTTCGTCAGTACTCTAATTTCCTATCTGACAACATCGAT | SmcR F75Y |
| JCV1306 | ATCGATGTTGTCAGATAGGAAATTAGAGTACTGACGAACGACATGATTAAGCACTT | SmcR F75Y |
| JCV1307 | TTCACGCTAAAGAAAACATCGCCAACCTGACCAACGCAATGATTGAGCTTGTGGTG | SmcR I96L |
| JCV1308 | CACCACAAGCTCAATCATTGCGTTGGTCAGGTTGGCGATGTTTTCTTTAGCGTGAA | SmcR I96L |
| JCV1309 | CCACCAACCGTACTAATCAGTTGCTGATCCAAAACATGTTCATCAAAGCCATTGAA | SmcR V140I |
| JCV1310 | TTCAATGGCTTTGATGAACATGTTTTGGATCAGCAACTGATTAGTACGGTTGGTGG | SmcR V140I |
| JCV1311 | ATTTGGCAAACCTGTTCCACGGCATTTTCTACTCGCTGTTTGTTCAAGCAAACCGC | SmcR C170F |
| JCV1312 | GCGGTTTGCTTGAACAAACAGCGAGTAGAAAATGCCGTGGAACAGGTTTGCCAAAT | SmcR C170F |
| JCV1313 | GTGCTGAATTTTGTGGTTCGTCAGTTTTCCAACTTCTTGACCGATCACATCG | HapR Y76F |
| JCV1314 | CGATGTGATCGGTCAAGAAGTTGGAAAACTGACGAACCACAAAATTCAGCAC | HapR Y76F |
| JCV1315 | TGGATGTGAAAACCAACCTACAAACTATCTGCAAAGAGATGGTGAAATTGGC | HapR L97I |
| JCV1316 | GCCAATTTCACCATCTCTTTGCAGATAGTTTGTAGGTTGGTTTTCACATCCA | HapR L97I |
| JCV1317 | CACCAACCGAACTAACCAACTGCTGGTAAGAAACATGTTTATGAAAGCGATG | HapR I141V |
| JCV1318 | CATCGCTTTCATAAACATGTTTCTTACCAGCAGTTGGTTAGTTCGGTTGGTG | HapR I141V |
| JCV1319 | ATGGCCAGCCTGTTCCACGGCATCTGTTACTCCATCTTCTTACAAGTGAACC | HapR F171C |
| JCV1320 | GGTTCACTTGTAAGAAGATGGAGTAACAGATGCCGTGGAACAGGCTGGCCAT | HapR F171C |
| AB324 | ATAAATCAAATTTGTAACTTTATTTCTGAGATGACTGTCCCATTTATCTTATTGATAAATCTGCGTAAAAAAGAAAAAGCACAATTTTATTACTAGTGTTT | $P_{vvpE}$ smcR, CAP, IHF BSs +25 bp left +12 bp right, F |
| AB325 | AAACACTAGTAATAAAATTGTGCTTTTTCTTTTTTACGCAGATTTATCAATAAGATAAATGGGACAGTCATCTCAGAAATAAAGTTACAAATTTGATTTAT | $P_{vvpE}$ smcR, CAP, IHF BSs +25 bp left +12 bp right, R |
| JDN54 | TAATGTTATTTGTAACACTATAAATAAGTAA | $P_{luxC}$ Site B, R |
| JDN55 | TTACTTATTTATAGTGTTACAAATAACATTA | $P_{luxC}$ Site B, F |
| JDN101 | GGATCCTCTAGAGTCGACCT | pMMB amplify backbone, F |
| JDN102 | GAATTCTGTTTCCTGTGTGAAATTG | pMMB amplify backbone, R |
| JDN103 | GATAACAATTTCACACAGGAAACAGAATTCATGGACTCAATCGCAAAGAGA | smcR in pMMB, F |
| JDN104 | GCATGCCTGCAGGTCGACTCTAGAGGATCCCTATTCGTGTTCGCGTTTATAGAT | smcR in pMMB, R |

**Table S5. Plasmids used in this study.**

| Name | Description | Reference |
|---|---|---|
| pAMS001 | hapR F171C Y76F in pJV387 | This study |
| pET28b | pET28b, 6X-his tag, $Kan^R$ | EMD |
| pJN08 | 6x-his-*smcR* in pET28b | [15] |
| pJN22 | $P_{tac}$-*smcR* (*V. vulnificus* ATCC 27562), in pMMB67EH-kanR | [15] |
| pJV064 | $P_{05222}$-mCherry, $P_{luxC}$-*gfp*; p15a origin, $CM^R$ | [16] |
| pJV387 | $P_{tac}$-*hapR* (*V. cholerae* E7946), in pMMB67EH-kanR | This study |
| pJV388 | $P_{tac}$-*luxR* (*V. campbellii* BB120), in pMMB67EH-kanR | This study |
| pJV389 | $P_{tac}$-*opaR* (*V. parahaemolyticus* RIMD 2210633), in pMMB67EH-kanR | This study |
| pJV390 | $P_{tac}$-*vcpR* (*V. coralliilyticus* OCN008), in pMMB67EH-kanR | This study |
| pJV391 | hapR Y76F in pJV387 | This study |
| pJV392 | hapR L97I in pJV387 | This study |
| pJV393 | hapR I131V in pJV387 | This study |
| pJV394 | hapR F171C in pJV387 | This study |
| pJV395 | SmcR F75Y in pMMB67EH-kanR | This study |
| pJV396 | SmcR I96L in pMMB67EH-kanR | This study |
| pJV397 | SmcR V140I in pMMB67EH-kanR | This study |
| pJV398 | SmcR C170F in pMMB67EH-kanR | This study |
| pJV79 | *luxR* in pET28b | [17] |
| pKM699 | *V. campbellii* $P_{luxR}$-*luxR* region (2.3 kbp fragment) in pLAFR2, $Tet^R$ | [18] |
| pLAFR2 | Empty vector cosmid, $Tet^R$ | [18] |
| pMMB67EH-kanR | Empty vector control; *kan*$^R$ | [11] |
